# Higher fitness yeast genotypes are less robust to deleterious mutations

**DOI:** 10.1101/675314

**Authors:** Milo S. Johnson, Alena Martsul, Sergey Kryazhimskiy, Michael M. Desai

**Affiliations:** Department of Organismic and Evolutionary Biology, Harvard University, Cambridge MA 02138; Quantitative Biology Initiative, Harvard University, Cambridge MA 02138; NSF-Simons Center for Mathematical and Statistical Analysis of Biology, Harvard University, Cambridge MA 02138; Division of Biological Sciences, University of California San Diego, La Jolla CA 92093; Department of Physics, Harvard University, Cambridge MA 02138

## Abstract

Natural selection drives populations towards higher fitness, but second-order selection for adaptability and mutational robustness can also influence the dynamics of adaptation. In many microbial systems, diminishing returns epistasis contributes to a tendency for more-fit genotypes to be less adaptable, but no analogous patterns for robustness are known. To understand how robustness varies across genotypes, we measure the fitness effects of hundreds of individual insertion mutations in a panel of yeast strains. We find that more-fit strains are less robust: they have distributions of fitness effects (DFEs) with lower mean and higher variance. These shifts in the DFE arise because many mutations have more strongly deleterious effects in faster-growing strains. This negative correlation between fitness and robustness implies that second-order selection for robustness will tend to conflict with first-order selection for fitness.

---

The dynamics and outcomes of adaptive evolution depend on the genetic variation available to a population. Because mutations interact epistatically, the availability and strength of beneficial and deleterious mutations can vary across genotypes (*1*). As “first-order” natural selection drives populations uphill in the fitness landscape, “second-order” selection can steer populations into regions with better uphill prospects (i.e. are more adaptable) or into “flatter” regions with fewer or less steep downhill paths (i.e. are more robust to deleterious mutations) (*2*–*5*). While there are several reports of second-order selection affecting evolutionary dynamics in microbes, viruses, and digital organisms (*4*–*8*), we still lack a general understanding of how and why genotypes differ in their robustness and adaptability.

Several recent studies have identified a consistent pattern of declining adaptability in laboratory microbial populations: less-fit genotypes tend to adapt more rapidly than more-fit genotypes (*9*–*13*). These differences in adaptability are at least partially explained by “diminishing returns” epistasis, in which individual beneficial mutations become less beneficial in more-fit genetic backgrounds (*10, 11, 14, 15*). In contrast to adaptability, genetic variation in mutational robustness has not been systematically characterized in any microbial system, and no analogs to diminishing returns or the rule of declining adaptability are known. There is evidence for both antagonistic and synergistic epistasis between pairs of deleterious mutations (*16*–*19*), but little is known about how the entire distribution of fitness effects of deleterious mutations changes across different genetic backgrounds.

There are several ways to define and measure mutational robustness (*2*–*4*). Here we consider only the single-step mutational neighborhood of a genotype, specifically focusing on the fitness effects of deleterious mutations. To understand genetic variation in single-step mutational robustness, we aim to measure how the distribution of fitness effects of deleterious mutations varies across genotypes, and how these differences arise from epistasis at the level of individual mutations. To do so, we would ideally like to measure the fitness effects of identical large sets of random deleterious mutations in multiple genotypes. To this end, we developed a high-throughput mutagenesis and fitness assay pipeline to measure the effects of sets of specific insertion mutations in a panel of *S. cerevisiae* genotypes (Fig. 1). We start from an existing yeast transposon mutagenesis library (*20*), which we down-sample to create a set of 1147 distinct plasmids (fig. S1). Each plasmid in this library contains a fragment of the yeast genome with a Tn7 transposon insertion. We then add random DNA barcodes to create a set of libraries that contain multiple copies of each plasmid, each tagged with a unique barcode. We sequence these plasmid libraries to associate each barcode with a specific insertion mutation and a specific library (Methods). We transform yeast strains with these libraries; homology-directed repair then creates the set of exactly identical transposon insertion mutants in each strain. While these insertion mutations are not a random sample of all naturally occurring mutations in yeast, they do represent an unbiased set of genomic disruptions (selected only to avoid artifacts, see Methods).

**Figure 1.**
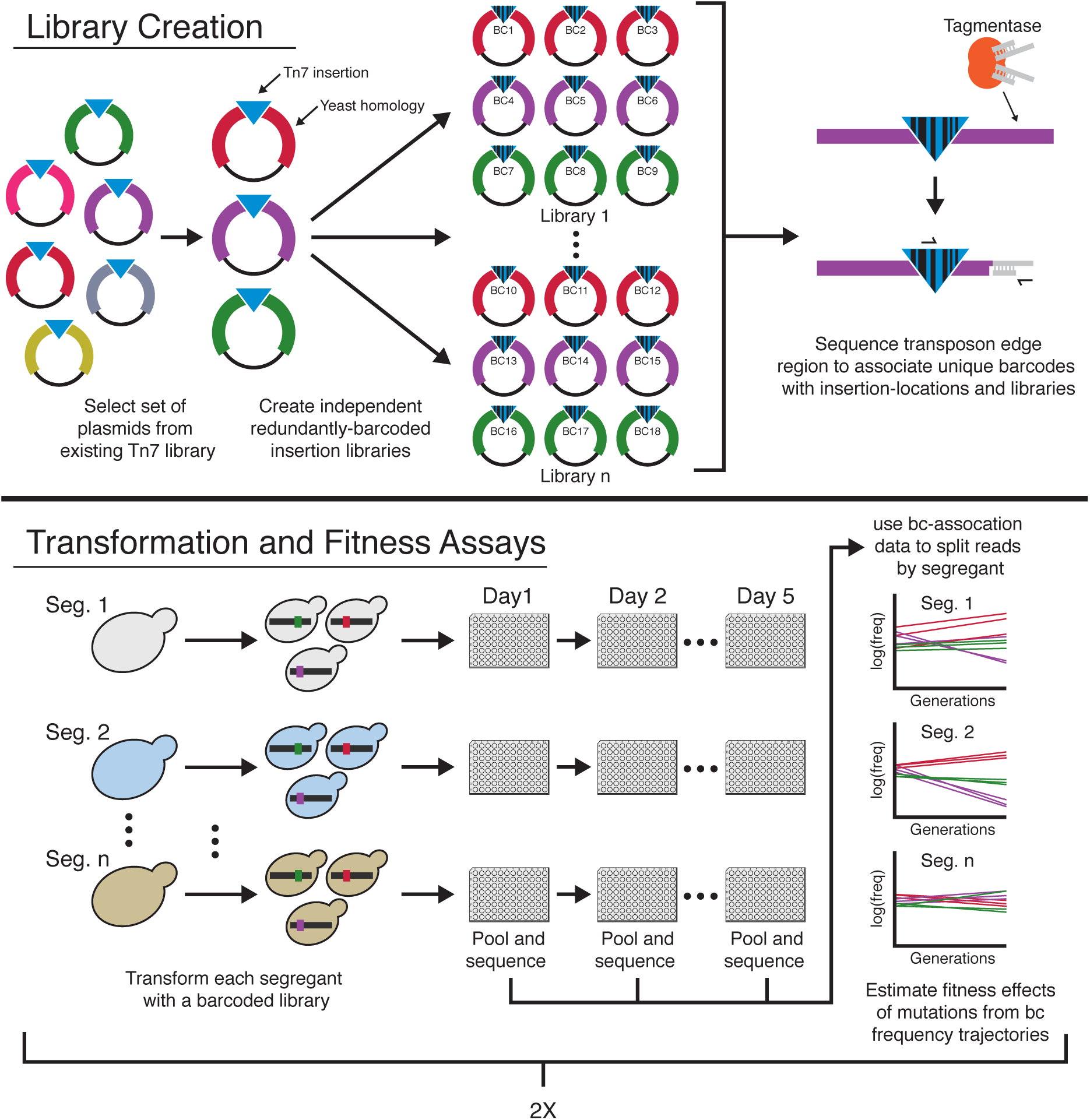
Schematic of mutagenesis and fitness assay pipeline. Plasmids with different colors indicate different regions of homology from the yeast genome.

To measure the fitness effect of each insertion mutation in each strain, we propagate the resulting mutant pools in batch culture for four growth cycles in our focal environment: rich media in 96-well microplate wells, with daily 1:2^10^ dilutions. We sequence the barcode locus at each transfer and estimate the fitness effects of mutations from barcode frequency trajectories. Because each insertion is represented by multiple uniquely barcoded transformants, we can identify and exclude outlier fitness measurements (likely due to transformation artifacts, multiple insertions, or pre-existing mutations; Methods, fig. S2). To estimate measurement noise, we conduct transformations and fitness assays in two replicates (Methods, fig. S3).

We applied this method to measure mutational robustness in F1 segregants derived from a yeast cross between a laboratory and a wine strain (BY and RM). These segregants differ at more than 35,000 loci and were sequenced and phenotyped in a previous study (*21*). In earlier work, we found that they span a 22.5% range of fitness in our focal environment, and we showed that segregants with lower initial fitness are more adaptable (*9*).

We first transformed 18 segregants selected at random from those studied in Ref. (*9*) (see Methods) with our 1147-mutation libraries. Due to gene essentiality, differences in cloning or transformation efficiency, or other complications, we were not able to measure the fitness effect of every mutation in every segregant (Methods); we successfully measured the effects of 710 mutations in at least one segregant, and an average of 414 per segregant. We found that 457 of these 710 insertion mutations (64.4%) have no detectable fitness effects in any of the segregants. Most of the remaining mutations were deleterious; the mean of the DFE is negative in all segregants. However, the distribution of fitness effects varied systematically across strains. Specifically, the mean fitness effect of all insertions decreases with the background fitness of the segregant (p=0.03, Fig. 2, A and B, fig. S4). In other words, more-fit segregants tend to be less robust with respect to random insertion mutations.

**Figure 2.**
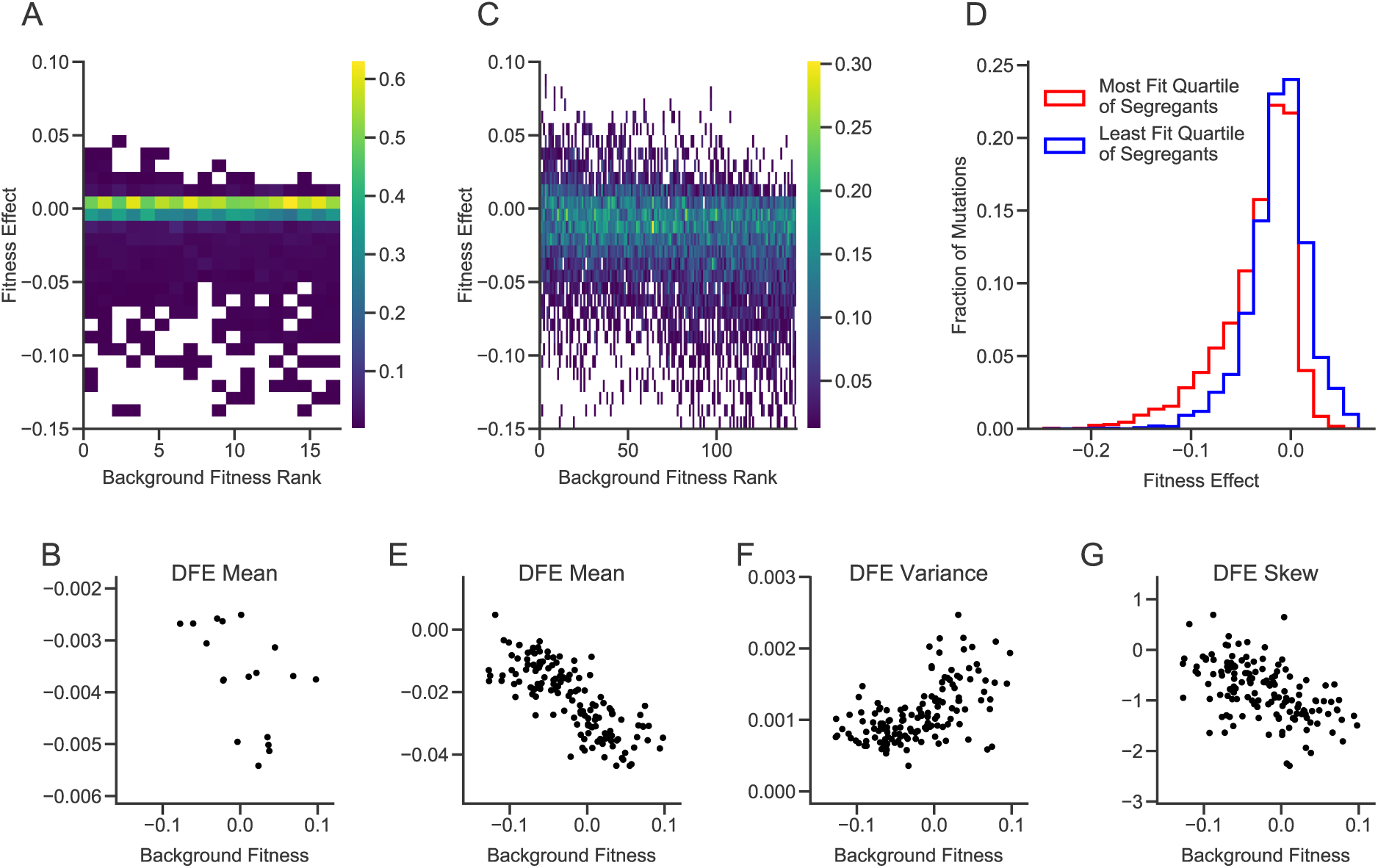
Distributions of fitness effects of insertion mutations. (**A)** Distributions of fitness effects in the large library experiment. Segregants are organized by background fitness. Color represents the fraction of mutations for each segregant in each fitness effect bin (see scale bar at right). (**B)** Relationship between background fitness and the mean of the DFE for the large library experiment. (**C)** Distributions of fitness effects in the small library experiment. (**D**) Combined distribution of fitness effects of most-fit and least-fit quartile of segregants in the small library experiment. (**E-G)** Relationship between background fitness and DFE statistics for the small library experiment.

Because we only measured fitness effects across 18 segregants and most mutations in our original library are indistinguishable from neutral in all segregants, we were not able to detect more subtle changes in the shape of the DFE or to connect DFE-level changes to patterns of epistasis for individual mutations. To address this limitation, we created a “small” library by selecting a subset of 91 insertions that had significant fitness effects in the largest number of segregants, and 5 intergenic insertions that appeared to be neutral in all segregants as negative controls (Methods). This small library filters out insertions with undetectable or rare effects but is otherwise unbiased. We transformed 163 segregants with this library and measured the fitness effect of each mutation in each segregant, as described above.

Again, we find that the mean of the DFE decreases with the fitness of the segregant (Fig. 2, C, D, and E). We also find significant correlations between background fitness and the variance and skew of the DFE, such that more-fit segregants have wider DFEs that are skewed towards more deleterious mutations (Fig. 2, C, F, and G, fig. S5, and fig. S6). These results imply that mutational robustness is negatively correlated with background fitness, so that second-order selection for these traits would be constrained by conflict with first-order selection for fitness.

The DFE is composed of individual mutations. To understand why the shape of the DFE varies between low and high fitness genotypes, we examined how the effects of these individual mutations vary. We observe a variety of patterns of epistasis (Fig. 3 and fig. S7), including nearly constant effects across backgrounds (e.g. PAH1, MME1) and diminishing returns (e.g. SIR3). However, the most frequent pattern is “increasing cost” epistasis, such that the same mutation is more deleterious in more-fit segregants (Fig. 3 and Fig. 4A). This is the deleterious-mutation analog of “diminishing returns” epistasis. Strikingly, this effect can cross zero: some mutations are beneficial in the least-fit segregants, neutral in intermediate-fitness segregants, and deleterious in higher-fitness segregants (e.g. RPL16A). However, this negative correlation is not universal, and a few mutations exhibit the opposite pattern (e.g. KRI1).

**Figure 3.**
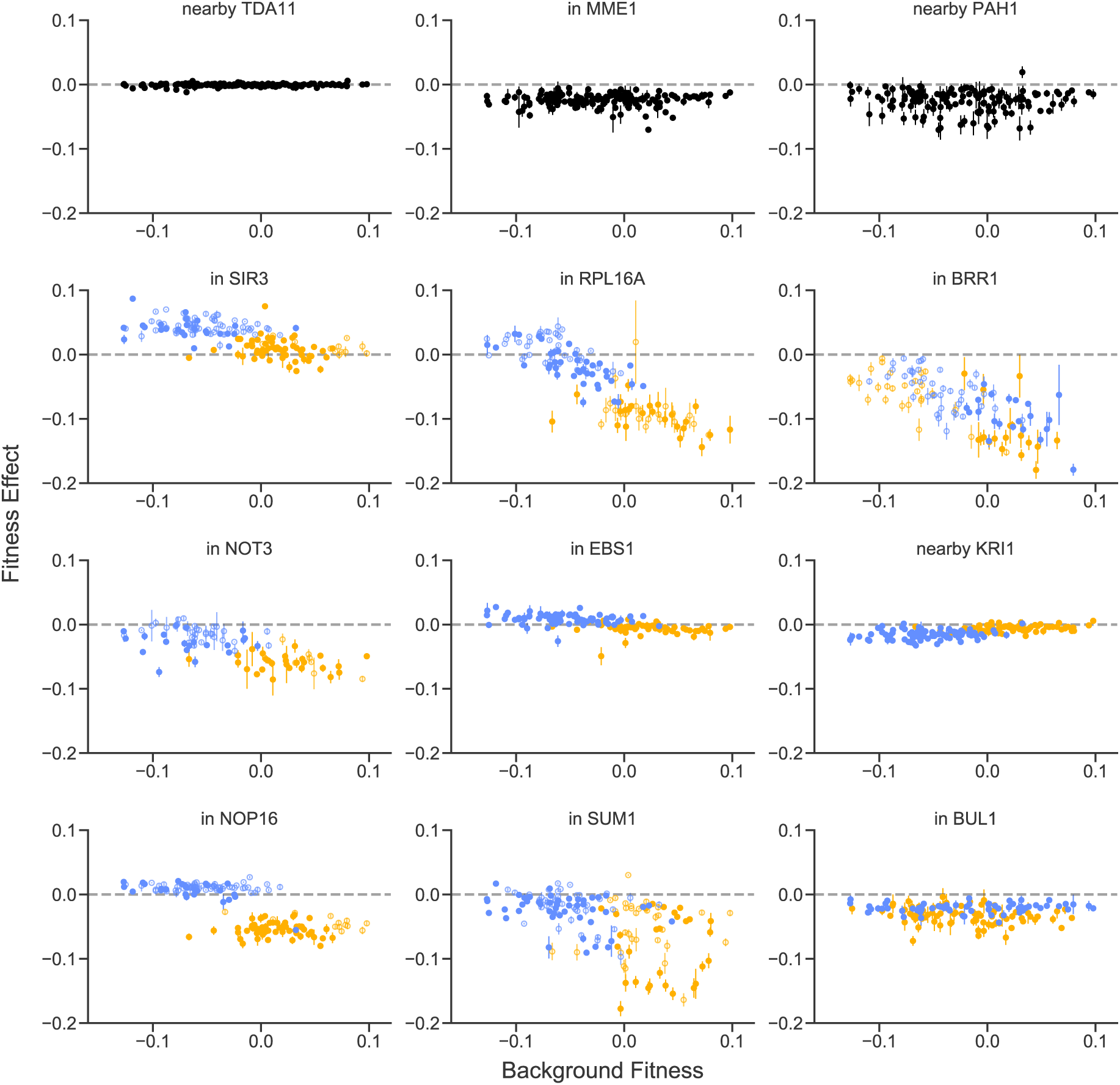
Patterns of epistasis for individual mutations. Fitness effects of 12 representative insertion mutations are plotted against segregant background fitness. The top left mutation is one of five putatively neutral insertions used as reference controls. Allelic state at the largest-effect quantitative trait locus for the fitness effect of each mutation is shown by yellow (BY) or blue (RM) color; allelic state at the second largest-effect quantitative trait locus is shown by closed (BY) or open (RM) symbol. Analogous plots for all insertion mutations are shown in fig. S8. Error bars represent standard errors (Methods).

**Figure 4.**
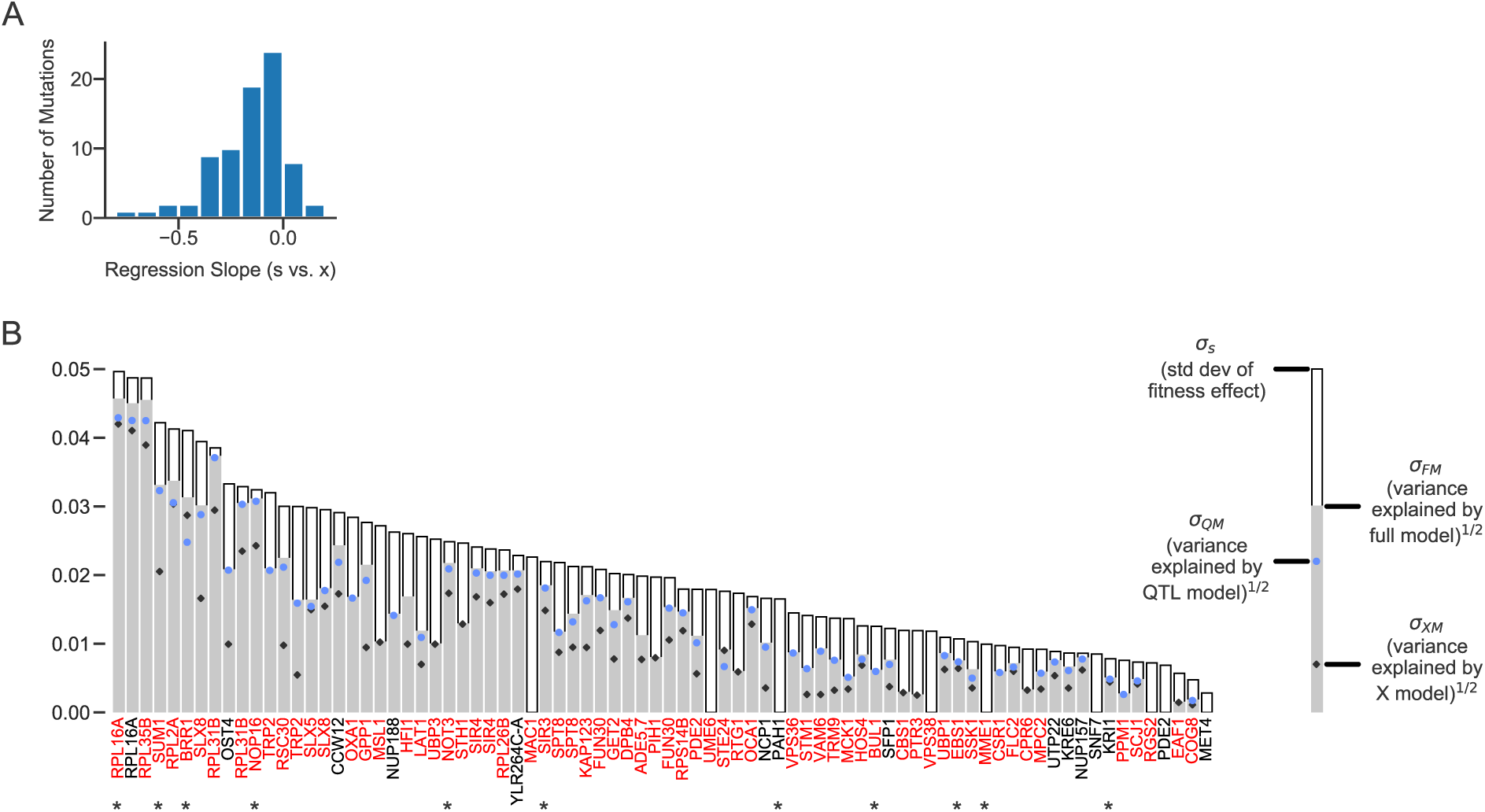
Genetic determinants of fitness effects. **(A)** Histogram of regression slopes between fitness effect and background fitness for each mutation. **(B)** For each mutation, the standard deviation of fitness effect across segregants and the square root of the variance explained by each of the three models (each model is only plotted if it is significant). Mutations shown in red or black are insertions in or near the corresponding gene respectively; stars indicate the mutations shown in Figure 3. Only mutations with fitness effect measurements in at least fifty segregants are shown.

In addition to the effect of background fitness, specific genetic loci can influence the fitness effect of a particular insertion mutation (e.g. NOP16). Because we have genotype information for each segregant, we can map the QTLs that affect the fitness effect of each insertion mutation (Methods, fig. S8 and fig. S9). We identified these QTLs using the same iterative procedure as in previous work (8, 18). For 62 out of 91 mutations, we detected at least one QTL that partially explains its fitness effect. To quantify how the initial fitness of the strain and the QTLs explain the variation in the fitness effect of each mutation across segregants, we fit three linear models to our data. The “fitness” model includes segregant fitness as the only predictor. The “QTL” model includes only the effect of segregant genotype at a small number of QTLs as predictors. The “full model” includes both segregant fitness and QTL effects as predictors.

We find that at least one of the three epistatic models is significantly better than the null model without epistasis for 71 of the 80 mutations for which we have measurements in at least 50 segregants. For mutations where initial fitness has explanatory power (64 cases, 80.0%), it explains on average 27.8% of variation in the fitness effect of a mutation. For mutations where QTLs have explanatory power (60 cases, 75.0%), they explain on average 48.3% of variation. Because many of the detected QTLs affect both segregant fitness and the fitness effect of the insertion mutation, QTL and fitness contributions are confounded. To understand which of the two predictors is more important, we use F-tests to compare the full model against the fitness and QTL models. We find that both effects are important for 53 (66.3%) mutations. The QTL model generally has more explanatory power than the background-fitness model, which is expected since it has more free parameters (8, 18). In many cases, though, the background fitness model explains close to as much variation as the QTL or full model, despite having only one parameter (Fig. 4B). This QTL analysis links the specific interactions between mutations to the broad pattern of diminishing returns/increasing costs epistasis we observe; QTLs that increase fitness also tend to undermine mutational robustness.

Because our assay environment involves batch culture with a 24-hour dilution cycle, faster growing strains spend more time in saturation phase. One potential explanation for the overall patterns of epistasis we observe is that insertion mutations affect survival during this saturation phase equally in all segregants, so more-fit strains are less robust because they spend longer in saturation. To investigate this possibility, we remeasured the fitness effect of each mutation in twelve segregants (chosen to span the full range of fitness of the 163 segregants, see Methods) on a finer timescale by sequencing barcodes and measuring cell density every 3-7 hours during growth. We found that both the differences in fitness between segregants and the fitness effects of insertion mutations arise almost exclusively during exponential growth (Methods, fig. S10, fig. S11 and fig. S12). Thus, changes in batch culture environments are not responsible for the patterns of epistasis we observe. Instead, our findings reflect differences in exponential growth rates: faster growing strains are more susceptible to deleterious mutations.

To search for functional explanations for the patterns of epistasis we observe, we classified insertion mutations into sets with particular epistatic patterns: those whose effects are influenced by a shared QTL, or those that had the strongest fitness-mediated epistasis (Methods). We looked for GO-term enrichment in the genes disrupted by the insertion mutations within each set, finding several enrichments at the 0.05 significance level (though none remain significant after multiple-hypothesis correction; Table S1). Most notably, the fitness-mediated epistasis set and the set associated with the most commonly-observed QTL (chromosome 14, between 360-420K) are enriched for ribosome and translation-related functions. This most commonly-observed QTL is also the strongest background fitness QTL, and includes variants in KRE33, a gene involved in small subunit assembly (discussed in more detail in (*9*)). Metabolic control theory provides a possible link between this functional information and the pattern of increasing cost epistasis in both of these sets of mutations: it predicts precisely this pattern between deleterious mutations in a linear metabolic pathway when fitness is correlated with flux through that pathway (*18, 22*). This theoretical work can extend beyond metabolism to any sequential biological pathway, and previous work has shown that deleterious perturbations of transcription and translation (due to either mutations or antibiotics) interact in the same way (*23*). While these results are not definitive, they suggest that further generalizations could be drawn from a deeper understanding of the cell-physiological basis of fitness-mediated epistasis.

The net effect of the variable patterns of epistasis we observe for individual mutations is a predictable change in the local properties of the fitness landscape. This “fitness landscape neighborhood” becomes less favorable in more-fit genotypes: uphill steps become flatter or even change to downhill, and many downhill paths become steeper. On such landscapes, first-order selection for high-fitness genotypes conflicts with second-order selection, and populations evolve towards more fit but less robust and less adaptable genotypes. Thus, populations may never attain true fitness peaks, instead reaching a dynamic balance between beneficial and deleterious mutations (*24*).

## Acknowledgements

We thank Elizabeth Jerison, Luke Rast, Alex Nguyen-Ba, Jeremy Yodh, Gregg Wildenberg, and members of the Desai lab for experimental assistance and/or comments on the manuscript. M.S.J. acknowledge support from the NSF Graduate Research Fellowship Program. S.K. acknowledges support from the BWF Career Award at Scientific Interface (Grant # 1010719.01), the Alfred P. Sloan Foundation (Grant FG-2017-9227), and the Hellman Foundation. M.M.D. acknowledges support from the Simons Foundation (Grant 376196), grant DEB-1655960 from the NSF, and grant GM104239 from the NIH. Computational work was performed on the Odyssey cluster supported by the Research Computing Group at Harvard University.

