## Supplemental Information for "Higher fitness yeast genotypes are less robust to deleterious mutations"

**This PDF file includes:**

Materials and Methods
Figs. S1 to S12
Tables S1 to S6
Data File Descriptions

**Other Supplementary Materials for this manuscript include the following:**

Data Files S1 to S3 (Excel format)

### Materials and Methods

#### Experimental Methods

##### Library creation

Our transposon mutagenesis library is derived from a previously existing diverse Tn7 plasmid library that includes insertions in ~300,000 locations in the yeast genome (*20*). In preliminary experiments, we used this library to transform yeast and found that a large percentage of successful transformants correspond to insertions in the ribosomal DNA (rDNA) array. To avoid this redundancy and to measure fitness effects accurately across many genetic backgrounds, we created a reduced plasmid library with ~1,000 non-rDNA insertions. To do so, we first cut the existing Tn7 library with I-PpoI (Promega), a restriction enzyme that cuts in the rDNA array, transformed this digested library into *E. coli* (strain DH10B (*25*)) and picked ~3,000 successful transformant colonies into single wells of 96-well plates. We then used Cartesian pooling-coordinate sequencing (*26*, *27*) to identify mutations to unique wells, using a modified TagMap method (*28*) in which we amplify the region spanning the edge of the Tn7 insertion (detailed below in "Modified TagMap Protocol"). We refer to the short regions of DNA next to the Tn7 insertion that are homologous to the yeast genome as "edges" here and in analysis code. By sequencing row, column, and plate pools, we identify the coordinates of those edges that are only present in a single well. We mapped slightly more than half of the edges to unique wells. Our ability to map edges to wells was coverage-limited, so additional sequencing likely would have allowed us to identify coordinates for more edges, but for our purposes this was sufficient. We aligned the edges to the yeast genome using bowtie2 (*29*). We record whether the Tn7 insertion site for each edge is in or near a gene / open reading frame (ORF). If the insertion is intergenic, we record the gene / ORF with the closest start site, up to 500 bp away. After eliminating edges that did not align uniquely to the genome, edges near the rDNA array (chr12 between 445K and 495K), and edges that were within 800 bp of a SalI restriction enzyme site (we use this enzyme during the next cloning step), we were left with a library of 1147 edges. We re-arrayed the corresponding set of *E. coli* clones, grew them to saturation, pooled them, and extracted plasmid DNA.

##### Modified TagMap protocol

To sequence the bases adjacent to the transposon edge, we used a "TagMap" protocol similar to Ref. (*28*), but we used the commercially available tagmentase (Illumina) instead of a specifically modified version, and used slightly different PCR and clean up protocols. We first tagmented plasmid DNA using the protocol from Ref. (*30*), scaled up 2X and with a higher starting DNA concentration (2.5-8 ng/ul). Next, we combined 5 μl tagmentation product, 22 μl 2X Kapa Hotstart Hifi MM, 8.8 μl 5 μM N7XX primer (Illumina), and 8.8 μl 10 μM TnR_to_P5_X primer, and ran the PCR protocol: 1) 95°C 5:00, 2) 98°C 0:10, 3) 63°C 0:30, 4) 72°C 0:25, GO TO step 2 12 times, 5) 72°C 5:00. We purified this PCR using AMPure XP magnetic beads (Beckman Coulter, 0.7X ratio), set up a second round PCR by combining 10 μl PCR 1 product, 25 μl 2X Kapa Hotstart Hifi MM, 7.5 μl 5 μM N7XX primer, and 7.5 μl 5 μM S5XX primer, and ran the same PCR protocol again. We purified this PCR using a two-sided bead size-selection (0.55X/0.65X). Note that our PCR steps are identical to Ref. (*30*) except that we skip the initial extension step.

##### Barcoding the library

To barcode this pooled plasmid library, we used Gibson assembly (*31*) to replace the interior of the Tn7 sequence in our initial library with an antibiotic resistance cassette and a random DNA barcode adjacent to the left arm of the transposon. First, we generated two intermediate plasmids, "pUC-Hyg" and "pUC-yNat" by Gibson assembling the pUC-19 backbone (https://www.addgene.org/50005/) and either the HygMX cassette (from FRP-1642 (https://www.addgene.org/58442/)) (*32*) or the yNatMX cassette (from pYTK078, (https://www.addgene.org/65185/)) (*33*). Note that we made these intermediate plasmids with both the ampicillin gene bla (beta-lactamase) along with the HygMX or yNatMX cassette in the transposon interior because of a previous barcode sequencing scheme; the libraries could be remade only using selection in *E. coli* on Hygromycin or clonNat (and Kanamycin for the existing library backbone); beta-lactamase is only included for historical reasons.

In order to create barcoded Tn7 libraries, we used Gibson assembly to combine SalI-HF-cut plasmids from our 1147-mutation plasmid library with purified PCRs of beta-lactamase and the HygMX or yNatMX cassette, from pUC-Hyg or pUC-yNat. These PCRs used primers tTEF_to_Tn7R, which adds homology to the Tn7 right edge, and Tn_BC_Connectorator_2, which adds the barcode sequence, a stop codon in each of three frames, a primer binding site, and homology to the Tn7 left edge. The stop codons in all three reading frames before the barcode are designed to ensure that the barcode sequence is never translated. We electroporated the purified product of this Gibson assembly into *E. coli* (strain DH10B).

For the large library experiment (hereafter "E1", "BT" in analysis code), we started with the library of 1147 mutations and barcoded the library separately 20 times, 10 each with the HygMX or yNatMX resistance cassette. The genomic location of all insertion mutations with measured fitness effects in E1 is shown in fig. S1. We used our modified TagMap method to sequence the transposon edges in each plasmid library, amplifying both the barcode region and the section of yeast genomic DNA next to the Tn7 edge. This allowed us to read barcodes and yeast genomic DNA sequences ("edges") together, so that we could associate barcodes both with specific mutations (edges) and plasmid libraries. For the small library experiment (hereafter "E2", "TP" in analysis code) we re-arrayed the 96 *E. coli* clones corresponding to the 96 selected mutations, grew them to saturation, pooled them, and extracted plasmid DNA. We then repeated the barcoding protocol on this pooled plasmid library, this time creating 96 libraries, 48 with the HygMX resistance cassette and 48 with the yNatMX resistance cassette. In both cases, each library (of the 20 or 96 libraries in E1 and E2 respectively) contains a unique set of barcodes, each of which is associated with a specific insertion mutation.

**Yeast strains**

The strains used in this experiment are segregants from a cross (previously described in Ref. *(21)*) between a prototrophic *MATa* BY parent and a prototrophic *MATα hoΔ::hphMX4 flo8Δ::natMX4 AMN1-BY* RM parent. For E1, we randomly selected 20 of these segregants that were used in a previous evolution experiment (*9*) and that did not have resistance to both clonNat and Hygromycin (one quarter of the segregants are doubly-resistant). Two segregants were excluded from analysis because they had a high percentage of outlier barcodes (see "Estimating fitness effects from barcode counts"). In E2, we used all 176 segregants from (*9*) that are not doubly-resistant. Note that our nomenclature for each segregant follows the convention in Ref. (*9*). Eight segregants were excluded from analysis because they had a high percentage of outlier barcodes (see "Estimating fitness effects from barcode counts"). The background fitness values we use throughout our analysis were measured in Ref. (*9*) using competitive fitness assays with a fluorescently-labeled reference strain.

**Yeast transformations**

Our yeast transformation protocol followed the method described in Ref. (*34*), with some minor modifications. Generally, we grew strains from freezer stocks overnight, diluted them 1:25 into YPD + Ampicillin (100 μg/mL) grew for 4 hours, washed the cells twice with sterile water, then resuspended in transformation mix and plasmid DNA cut with NotI-HF (cut at 37°C for 3 hours, then heat inactivated at 65°C for 10 min). We heat shocked this mixture at 42°C for 1 hour, recovered in YPD + Ampicillin for 2 hours, and selected in liquid YPD supplemented with the appropriate antibiotic (Hygromycin at 300 μg/mL and/or clonNat at 20 μg/mL). For most transformations, pre-transformation growth was carried out in 5mL of media in test tubes on a roller drum, recovery was in 1-2 mL of media in a test tube on the roller drum, and selection for successful transformants was carried out in 25 mL of media in baffled flasks on a standing shaker. All growth was conducted at 30°C. After antibiotic selection (~36 hours), we made frozen glycerol stocks of the transformed libraries. In E2, the transformations corresponding to replicate 1 were done in 96-well format using the BiomekFX pipetting robot (Beckman Coulter), but the transformations for the second replicate were done in tubes as per the protocol described in Ref. (*34*); we found the 96-well transformations had much lower efficiency than the tube-based transformations. Segregants with Hygromycin resistance were transformed with yNatMX libraries, those with clonNat resistance were transformed with HygMX libraries, and those with neither resistance could be transformed with either. Within these constraints, plasmid libraries were randomly assigned to segregants. All singly-resistant segregants were selected on both Hygromycin and clonNat to avoid cassette replacement during transformation.

**Bulk fitness assays**

All bulk fitness assays (BFAs) were performed in YPD (1% Bacto yeast extract (VWR #90000-726), 2% Bacto peptone (VWR #90000–368), 2% dextrose (VWR #90000–904)) in unshaken flat-bottom polypropylene 96-well plates at 30°C. All 96-well liquid handling was performed on a BiomekFX robot (Beckman Coulter). In E1, we diluted 1 ml of frozen stocks into 18.5 ml YPD + antibiotics and distributed 128 μl of this mixture into 96 wells of a 96-well microplate. In E2, we diluted 128 μl of frozen stocks into 1.92 ml YPD + antibiotics, split between 16 wells (8 μl of frozen stock into 120 μl YPD per well). We allowed these initial cultures to grow to saturation (~36 hours), then completed the first transfer, marking timepoint zero. After this initial grow-up, we grew yeast in 128 μl YPD + Ampicillin per well, with daily $1:2^{10}$ dilutions for 4 cycles (40 generations). In E1, each assay took place in all 96 wells of a 96-well microplate, and all wells from each plate were mixed together during the $1:2^{10}$ dilution. We mix wells during the transfer step to increase the population size of barcode lineages and thereby reduce noise in their frequencies. In E2, the mutant population descended from each segregant was distributed across 16 wells and those 16 wells were mixed together during the $1:2^{10}$ dilution. Logistically, each population occupied the well with the same coordinates in 16 separate 96-well plates (e.g. segregant 1 was in well B7 in all plates); because there were 176 segregant populations in E2, we maintained them in a total of 32 96-well plates. During E1, we pooled together equal volumes from all plates, pelleted cells, and froze pellets at -20°C. During E2, since there were 176 segregants but only 96 plasmid libraries, we made two pools at each timepoint, pooling equal volumes from all wells of 16 of the 32 96-well microplates for each, pelleted cells, and froze pellets at -20°C. We are able to pool the cultures at this stage because we know which barcodes correspond to which plasmid library, and therefore which segregant-assay, so we can divide the data during sequencing analysis. For both experiments, the second replicate was done on different days with a different batch of YPD. We excluded data from the first timepoint (T0) of experiment 2 replicate 1 because of pipetting errors during the pooling step.

**Sequencing library preparation**

We used the Yeastar Genomic DNA Kit Protocol I (Zymo Research) to extract DNA from ~1.5 ml of pelleted yeast culture. For E1 (both replicates) and E2 replicate 2 we used 2 DNA extractions for each timepoint, while for E2 replicate 1 we used only one DNA extraction because we needed less coverage for a smaller number of barcodes. To produce Illumina-ready, dual-indexed, UMI-tagged (Unique Molecular Index) fragments, we use a two-step PCR protocol similar to the one used in Ref. (*35*). For each of the timepoint-samples for E1 (both replicates) and E2 replicate 2, we performed 4 50 μl first round reactions. For E2 replicate 1 and the growth curve experiment, we performed 2. Briefly, we first combined 19 μl gDNA, 25 μl 2X Kapa Hotstart Hifi MM, 3 μl 10 μM TnRS1 primer, and 3 μl 10 μM Tn*FX* primer, and ran the PCR protocol: 1) 95°C 3:00, 2) 98°C 0:20, 3) 60°C 0:30, 4) 72°C 0:30, GO TO step 2 2 times, 5) 72°C 1:00. We purified this PCR with AMPure XP magnetic beads (0.85X ratio). We set up a second reaction by combining 25 μl purified PCR 1 product, 1.5 μl dd$H_2O$, 10 μl Kapa Hifi Buffer, 1 μl KAPA HiFi HotStart DNA Polymerase, 5 μl 5 μM N7XX primer (Nextera), and 5 μl 5 μM S5XX primer (Nextera), and ran the PCR protocol: 1) 95°C 3:00, 2) 98°C 0:20, 3) 61°C 0:30, 4) 72°C

0:30, GO TO step 2 17 times, 5) 72°C 2:00. We purified this PCR with AMPure XP magnetic beads (0.85X ratio). For some libraries, we repeated this final clean-up step to remove any remaining small fragments. We standardized library concentration and checked size distributions using a Qubit fluorometer (ThermoFisher) and Tapestation (Agilent). We sequenced these final libraries with single-end 75bp reads on a NextSeq (Illumina).

#### Data Analysis Methods

**Associating barcodes with specific mutations**
To find barcodes in our barcode association sequencing reads, we sequentially use a list of increasingly less strict regular expressions (using the python regex module https://pypi.org/project/regex/):
```
'(GTAGAA)(\D{28})(GCGTCA)',
'(GTAGAA)(\D{26,30})(GCGTCA)',
'(GTAGAA){e<=1}(\D{28})(GCGTCA){e<=1}',
'(GTAGAA){e<=1}(\D{26,30})(GCGTCA){e<=1}'
```
To find the edge sequence (the nucleotides adjacent to the transposon, which are used to identify the location of the insertion), we first look for the Tn7 sequence that marks the end of the transposon ('TCCGCCCACA'), allowing for one error if there is no exact match. In E1, we define the "edge" as the first 30 bp after this match. Since we already know the limited set of mutations in E2, we use only the first 15 bp, which allows us to use reads where tagmentation occurred between 15 and 30 bp from the transposon. If we can extract both the barcode and edge and do not detect the tagmentase (Tn5) sequence in the 15 or 30bp of the edge sequence we are trying to map, we add a count for that combination.

After counting barcode-edge combinations, we error correct barcodes and edges using a deletion-based algorithm that allows for sequences with up to 3 errors to be corrected (see "Deletion-error-correction algorithm" below). We filter out barcodes that are associated with more than one edge, then assign barcodes to a plasmid library if more than 95% of their reads are from that library and they have more than 3 reads total. For E2, we used Cartesian pooling-coordinate sequencing, so we assign barcodes to a plasmid library only if we can successfully assign them to a row and column using the same rules (e.g. >95% of row-library reads are from row A, >3 row-library reads).

**Deletion-error-correction algorithm**
This algorithm takes advantage of the fact that all errors should be connected by single insertions, deletions, and substitutions by using deletion neighborhoods to speed up the error-correction process. The algorithm uses as input a list of barcodes and total counts (reads) corresponding to each barcode. The steps are listed below:
1) Make deletion neighborhoods
    a) For each observed sequence, create the set of single-base deletions at each position
    b) Connect sequences with overlapping single-base deletion sets
        i) Note: this overlap indicates that the two sequences are separated by one "edit": a substitution, insertion, or deletion
2) Within each neighborhood, define the error-free sequences we are looking for ("peaks" in analysis code) by these criteria:
    a) Sequence does not contain an uncalled base ("N")
    b) Sequence has no single-edit neighbors with more total counts
    c) Sequence has more than (3 for barcodes, 20 for edges) total counts
    d) Sequence is more than 3 edits away from any peak with more total counts
3) Within each neighborhood, error correct non-peak sequences:
    a) Check the Levenshtein (edit) distance between each non-peak sequence and each peak sequence. If the edit distance is less than or equal to 3, the sequence error-corrects to the peak

sequence. If a sequence is within 3 edits of more than one peak, it corrects to the sequence with higher total counts

b) This step uses the python Levenshtein module (https://pypi.python.org/pypi/python-Levenshtein/0.12.0#license)

**Determining barcode counts**

For each read in our BFA sequencing, we first check that the inline index (first 8 bp after the unique molecular index (UMI)) is what we expect, and that the average quality score of the barcode region is at least 25. Then we use the same set of regular expressions as above (but with reverse complements of the sequences), to extract barcodes. 91-97% of reads pass these filters and yield barcodes that we count. We then combine barcode counts from separate sequencing libraries that correspond to different timepoints for the same assay.

We record the number of UMIs associated with each barcode as well as its raw counts, but since the UMIs are only 7 bases long, we expect a significant number of repeats by chance, so we simply use the raw counts. In preliminary work we used the UMI data to identify when libraries had experienced significant bottlenecks during the first round of PCR, which can lead to noisy read-count data. In our final analysis we ignore UMI information.

To assign counts to known barcodes, which are associated with edges and plasmid libraries, we use a simplified version of the deletion-error-correction method. First, we generate the single-bp deletion neighborhood for every known barcode (from barcode-association sequencing). For each barcode from the BFA read data, we then check if it is a known barcode. If it is not, we check if its single-bp deletion neighborhood overlaps with any known barcode (if it is a single error away from a known barcode), and if so, add a count for that barcode. We are able to match our observed barcodes to known barcodes for 92-96% of reads that pass the quality and regex filters described above. Any barcodes that match two known barcodes using this method are excluded (this is very rare, but does happen for ~0.01% of barcodes). The barcode association also tells us which plasmid library, and therefore which assay/segregant, this count belongs to, so we split counts based on these associations. This procedure results in separate barcode count files for each assay/segregant.

**Estimating fitness effects from barcode counts**

After excluding timepoints with less 5,000 counts, we measure the log-frequency slope for each barcode between each pair of consecutive timepoints, excluding timepoints where the barcode has less than 10 counts. For E1, we scale these fitness values, $s$, by the median log-frequency slope of all barcodes at each time interval. Since most mutations in this experiment are neutral, this corrects for differences in mean fitness between timepoints. In E2, we use the median log-frequency slope of the set of barcodes corresponding to the five putatively neutral mutations to make the mean fitness correction. We exclude assays with less than 5 neutral barcodes or less than 3 usable timepoints. We then calculate barcode fitness effects by averaging these scaled values over time intervals in the assay.

In preliminary analysis, we observed a variable but sometimes significant number of barcodes with fitness effects that deviated significantly from most other barcodes associated with the same mutation. This effect was most evident when the deviation was towards a more beneficial fitness effect, when high read counts in late timepoints can rule out the possibility of deviations due to noise. These deviations may be due to transformation artifacts (including two mutations being transformed together) or pre-existing mutations in the transformed population. Our redundant barcoding system allows us to identify and exclude these barcodes. Specifically, we use a log-likelihood ratio test to identify barcodes with a fitness effect significantly different from the median fitness effect observed among all barcodes corresponding to the same mutation (see "Excluding outlier barcodes" below). Because these types of artifacts are difficult to avoid, we suggest that high-throughput studies of this type should make redundant measurements and exclude outliers whenever possible. After excluding outlier barcodes, we reduce the number of barcodes for each mutation in each assay to a maximum of 7 by randomly combining counts from individual barcodes into combined barcodes (cBCs). This reduces the per-barcode noise for assays with many barcodes, while still preserving 7 independent fitness measurements if possible. Finally, we repeat the fitness measurement protocol on these cBCs. For some assays, the rate of these outlier barcodes is too high to effectively exclude them, likely due to a mutation that arose before transformation; we identify these assays by visually inspecting the data and frequency trajectories and exclude them from further analysis (Table S2). We excluded 2 segregants from E1 and 8 segregants from E2 based on high rates of outlier barcodes.

We only use fitness measurements for mutations with at least 3 cBCs total in the two replicate fitness assays. For each replicate, we calculate the mean *s* across cBCs and the standard error of that measurement, which is the square root of the sum of the squared standard error for the mutation and the squared standard error of putatively neutral cBCs:

$$s_{rep} = \frac{\sum_{cbcs} s_{cbc}}{num\ cbcs},$$

$$SE_{rep} = \sqrt{SE_{(s_{cbc})} + SE_{(s_{neutral\ cbcs})}}.$$

For replicates with only one cBC, we use the standard deviation of *s* in the cBCs from the other replicate as an estimate of $SE_{(s_{cbc})}$. We average across replicates using inverse-variance-weighting (similar to the approach in (*36*)):

$$\bar{s} = \frac{\sum_{reps} \frac{s_{rep}}{SE_{rep}^2}}{\sum_{reps} \frac{1}{SE_{rep}^2}}.$$

We calculate the standard error on this final *s* measurement based on the *s* values measured for each cBC:

$$SE_{\bar{s}} = \frac{\sqrt{\frac{\sum_{cbcs}(s_{cbc}-\bar{s})^2}{num\ cbcs-1}}}{\sqrt{num\ cbcs-1}}.$$

We chose to use this conservative measure of standard error instead of the one given by inverse-variance weighting because the inverse-variance weighted error does not capture biological error between the two replicates (if both replicates have low $SE_{rep}$, but $s_{rep}$ is very different between replicates, the error calculated using the inverse-variance weighting method will be low, but our measure of error will capture these differences).

Correlations between $s_{rep}$ for the two replicates are shown in fig. S3A (E1) and fig. S3B (E2). Correlations between $\bar{s}$ for the two experiments are shown in fig. S3C.

To test whether mutations have effects significantly different from zero, we fit all cBC *s* values for a single mutation by ordinary least squares with an independent variable for replicate according to the equation: $s_{cbc} = s_{mut} + B_{mut}r + e_{mut}$, where $e_{mut}$ is a normally distributed noise term, $r$ is an indicator variable for replicate, $B_{mut}$ is the (fitted) coefficient representing differences between replicates, and $s_{mut}$ is the intercept term which represents the fitness effect of the mutation. We use the t-statistic for the intercept to calculate p-values for whether $s_{mut} \neq 0$ for each mutation.

**Excluding outlier barcodes**

To test for outlier barcodes, for each mutation with at least 3 barcodes (if a mutation has less than 3 barcodes no exclusions are possible), we pool counts from all barcodes with a measured *s* within 0.01 of the median

$s$ for that mutation. We call this set of pooled-counts "the within-mutation neutral reference" ("WMNR"). For a barcode where $s$ is more than 0.01 different than the median $s$, we consider only that barcode's counts and the WNMR's counts. We use a model that assumes deterministic changes in frequency and multinomial reads, where the log-likelihood of the data at each timepoint $t$, given a frequency prediction $f_{k,t}$ and number of reads $n_{k,t}$ for each of $k$ lineages, is $LL = Constant + \sum_t \sum_k n_{k,t} \log(f_{k,t})$. We find the log-likelihood of two hypotheses: 1) The barcode and the reference have the same fitness (one free parameter corresponding to relative starting frequency) and 2) The barcode and the reference have different fitnesses (two free parameters, relative starting frequency and relative fitness). Finally, we exclude barcodes with a log-likelihood ratio above a heuristic cutoff (We use 15, see below).

While the deterministic-frequencies-multinomial-reads model is not explicitly correct for these experiments, we found using simulated data that this log-likelihood ratio can effectively eliminate barcodes with a fitness effect more than 0.025 different from the mutation fitness, with more sensitivity when the deviation is toward a more beneficial effect. Specifically, we simulated bulk competition experiments with parameters (for DFE mean, std. deviation, and read depth) estimated from the E1 or E2 experiment, and a random percentage of outliers (uniform from 0-10%), whose fitness effects are drawn from a uniform distribution from -0.1 to 0.1. Fig. S2A the percentages of outliers that are correctly excluded and the percentage of false positives at various log-likelihood ratio cutoffs. Based on these results, we chose 15 as a conservative log-likelihood ratio cutoff. Examples of outliers excluded using this method are shown in fig. S2 B and C.

**Missing fitness measurements**

Because we did not measure the fitness effect of every mutation in every strain, it is possible that our DFE-level results are skewed by missing data. To test whether this is the case, we looked at largely beneficial or deleterious mutations and asked whether the number of these mutations successfully measured in each segregant varies as a function of background fitness. We defined these sets of mutations as having positive or negative fitness effects on average (across segregants) and having a statistically significant fitness effect in at least 2 segregants. The only significant regression was between background fitness and the number of deleterious mutations measured in E2, such that deleterious mutations were less likely to be measured in more fit backgrounds (fig. S6). This small effect suggests that we may have systematically missed some measurements of deleterious mutations in fit segregants, which strengthens our conclusions about background fitness effects on the DFE. Data for individual mutations (Fig. 3 in the main text) also support our conclusion that the DFE-level patterns we see are not caused by missing data.

**Modeling genetic determinants of fitness effects**

For each mutation, we perform a Benjamini-Hochberg correction at the 0.05 level on the p-values from each segregant to determine in how many segregants it has a significant effect. Next, if the mutation has fitness effect measurements in at least 10 segregants, we fit a series of models to the fitness effect data for each mutation:

Parameters:

| | |
|---|---|
| $s_j$ | (Measured) fitness effect in segregant $j$ |
| $x_j$ | (Measured) background fitness of segregant $j$ |
| $g_{ij}$ | (Measured) segregant $j$'s genotype at locus $i$ |
| $A$ | (Fitted) intercept term |
| $B$ | (Fitted) background fitness coefficient |
| $Q_i$ | (Fitted) locus $i$ coefficient |
| $e_j$ | normally distributed noise parameter |

Background-fitness model (model (i)):

$$s_j = A + Bx_j + e_j$$

QTL model (model (ii)):

$$s_j = A + \sum_i Q_i g_{ij} + e_j$$

Full model (model (iii)):

$$s_j = A + Bx_j + \sum_i Q_i g_{ij} + e_j$$

To find QTLs and confidence intervals, we use the same iterative approach applied in (*9*) and (*21*). We use the resulting list of QTLs for the QTL model. We also create an extended list of QTLs by adding any new QTLs that explain the residuals from the background-fitness model. We use this extended QTL list for the full model. We use the python package *statsmodels* to fit each of these models by ordinary least squares. For mutations in E2, we subsample our data, choosing half of the segregants (no replacement) 1,000 times to calculate a 95% confidence interval for the $R^2$ values for each model.

**Modeling genetic determinants of DFE statistics**
For each segregant, we calculate statistics (mean, median, variance, skew, kurtosis) of the distribution of fitness effects using all "measured" mutations (those with data from at least 3 barcodes across two replicates) (Data File S1). We determine the number of significantly beneficial or deleterious mutations by performing a Benjamini-Hochberg correction at the 0.05 level on the p-values from all measured mutations in the segregant in question. Next, we fit the same set of models as above, this time to each of the statistics (including the number of beneficial and deleterious mutations). We subsample our data, choosing half of the segregants (no replacement) 1,000 times to calculate 95% confidence intervals for each statistic. Fig. S4 (E1) and fig. S5 (E2) show the relationship between these DFE statistics and background fitness.

**Multi-hit QTLs and GO-Term enrichment**
The locations and effect sizes of all detected QTLs are listed in Data File S2. We plotted the location of all QTLs detected in our experiment (fig. S8) and identified 8 "multi-hit QTL regions" as those where more than two mutations had detected QTLs. Five of these multi-hit QTL regions overlapped with QTLs that had effects on fitness in YPD in Ref. (*9*) and/or Ref. (*21*) (Data File S2).

To quantify if the mutations interacting with each multi-hit QTL had any functional connections, we performed GO-Term Enrichment analysis using the GOATOOLS python library (*37*). We tested the group of genes corresponding to mutations in each multi-hit QTL group, and we also tested the group of genes corresponding to the mutations with the largest background fitness coefficients in the full model (the mutations with the strongest background-fitness effects). We used the set of genes corresponding to all mutations for which we measured a fitness effect in at least 10 segregants as the background population for this testing. After correcting for multiple hypotheses using the Benjamini-Hochberg method, we found no significant GO-Term Enrichment. The enrichments that were significant before the multiple hypothesis testing correction are listed in Table S1.

Two of the multi-hit QTLs are near the drug marker replacements in the parental strains (chr04 40000-70000, near *hoΔ::*hphMX4, and chr05 360000-400000, near *flo8*Δ::natMX4). Fig. S9 shows the relationship between fitness effect and background fitness for mutations with detected QTL effects in these regions, with points colored according to segregants' genotypes at the loci in question. While several of these mutations clearly have negative correlations between fitness effects and background fitness, these QTLs do not have a large effect on background fitness and therefore do not appear to drive these negative correlations (in contrast to other QTLs detected in our study, e.g. see NOP16 in Fig. 3 of the main text). If we exclude this set of mutations from our analysis our conclusions are generally unchanged, although the correlation between background fitness and the mean of the DFE is no longer significant at the 0.05 level in E1 (p=0.0504).

**Choosing mutations for the second experiment**

To select 91 mutations with significant fitness effects for our E2 experiment, we used a different fitness effect estimation procedure than described above. Specifically, after inferring selection coefficients for each barcode as described above, we modeled the selection coefficient $s_{imb}$ of barcode $b$ in strain $i$ that corresponds to mutation $m$ as

$$s_{imb} = \mu_{im} + \delta_{imb} + e_{imb},$$

where $\mu_{im}$ is the true fitness effect of mutation $m$ in strain $i$, $\delta_{imb}$ is the effect of transformation in the mutant carrying barcode $b$, and $e_{imb}$ is the measurement error. We assumed that all $\varepsilon_{imb}$ are independent, normally distributed random variables with mean 0 and variance $\sigma^2_{\mathrm{err}}$. We modeled the transformation artifact as a random variable that takes value 0 with probability $1–p_{\mathrm{tr}}$, and takes value $–\Delta_{imb}$ with probability $p_{\mathrm{tr}}$, where all $\Delta_{imb}$ are independent and normally distributed with mean $\mu_{\mathrm{tr}}$ and variance $\sigma^2_{\mathrm{tr}}$. Thus, the likelihood of observing the value $s_{imb}$ in this model is

$$L(s_{imb};\ \mu_{im}, \theta) = (1 - p_{\mathrm{tr}})N(s_{imb}; \mu_{im}, \sigma^2_{\mathrm{err}}) + p_{\mathrm{tr}}N(s_{imb}; \mu_{im} - \mu_{\mathrm{tr}}, \sigma^2_{\mathrm{err}} + \sigma^2_{\mathrm{tr}}), \qquad [1]$$

where $N(x; \mu, \sigma^2)$ is the normal probability density function with mean μ and variance $\sigma^2$ and $\theta = (p_{\mathrm{tr}}, \mu_{\mathrm{tr}}, \sigma^2_{\mathrm{tr}}, \sigma^2_{\mathrm{err}})$ is the vector of "global" parameters (i.e., those that are common to all mutations and strains). The likelihood of the entire data $\{s_{imb}\}$, which contains measurements of all barcode fitness effects in all strains, is given by

$$L(\{s_{imb}\};\ \{\mu_{im}\}, \theta) = \prod_i \prod_m \prod_b L(s_{imb};\ \mu_{im}, \theta). \qquad [2]$$

This model has four global parameters ($p_{\mathrm{tr}}$, $\mu_{\mathrm{tr}}$, $\sigma^2_{\mathrm{tr}}$, $\sigma^2_{\mathrm{err}}$) and thousands of parameters (the $\mu_{im}$) which represent the fitness effects of individual mutations in different genetic backgrounds. Due to this large number of parameters, maximizing the likelihood function [2] directly using standard methods is not feasible. We therefore maximized it using an iterative scheme, as follows. Given the estimated values of the global parameters $\theta^{(k)}$ at iteration $k$, we maximized the likelihood functions [1] independently for each mutation in each strain with respect to $\mu_{im}$, while holding $\theta^{(k)}$ constant. We thereby obtained an estimate $\mu^{(k+1)}_{im}$. We then maximized the likelihood function [2] with respect to $\theta$, while holding all $\mu^{(k+1)}_{im}$ constant. We thereby obtained an estimate $\theta^{(k+1)}_{im}$. We continued this procedure until the value of the likelihood function [2] and all estimates of global parameters $\theta$ converged.

This procedure yielded the following estimates of the global parameters:

$$\begin{aligned} \hat{p}_{\mathrm{tr}} &= 0.19, \\ \hat{\mu}_{\mathrm{tr}} &= 0.23, \\ \hat{\sigma}^2_{\mathrm{tr}} &= 5.23, \\ \hat{\sigma}^2_{\mathrm{err}} &= 0.16. \end{aligned}$$

The estimated values of the $\hat{\mu}_{im}$ are given in Data File S3. To decide whether the effect $\mu_{im}$ of mutation $m$ in strain $i$ was statistically different from 0, we used the likelihood ratio test. Specifically, we calculated the statistic

$$R = 2\big(\log L\big(s_{imb};\ \mu_{im}, \hat{\theta}\big) - \log L\big(s_{imb};\ 0, \hat{\theta}\big)\big),$$

and obtained its p-value from the $\chi^2$-distribution with 1 degree of freedom. Then, for each of the 18 segregants, we used the Benjamini-Hochberg procedure to obtain the list of mutations that had fitness effects significantly different from zero with false discovery rate of 0.001. The estimated p-values and mutations identified as significant are given in Data File S3.

Using this method, we selected mutations that had significant effects in at least six segregants and were identified during our initial Cartesian pooling-coordinate sequencing step (this was necessary in order to locate the corresponding *E. coli* strain). This yielded a set of 92 mutations. Three of these mutations were in the gene FUN30, so we excluded one of these mutations to bring the total to 91. We chose our five putatively neutral mutations from the set of mutations with no significant fitness effects and fitness effects measured within 0.5% of zero for all segregants.

#### Growth Curve Experiment

**Growth curve experiment methods**
We chose 12 strains from the E2 experiment that spanned a wide range of initial fitness values and had non-overlapping sets of barcodes across both replicates (Table S3). One segregant was excluded from analysis because cell density data showed that it was not at saturation prior to inoculation at the first timepoint. We inoculated the transformed libraries of both replicates for each strain using the same dilution as in the E2 experiment but organized the plates so that each replicate of each strain (24 populations) was in 8 wells of a 96-well plate (2 plates total). Thirty-four hours after inoculation, we diluted each of these plates 1:$2^{10}$ into 24 96-well plates (48 total plates). Additionally, we combined equal volumes from replicate 2 for each strain (a total of 12 populations) and diluted this mixture 1:$2^{10}$ into 1 liter of YPD + Ampicillin in a 4-liter baffled flask, which was shaken at 170 rpm at 30°C. Note that this flask experiment is the only time we mixed and grew different segregants together in this entire project.

At a series of chosen timepoints between 3 and 7 hours apart we destructively sampled the 48 plates and sampled volume from the flask (details on the numbers of plates and volumes sampled are listed in Table S4). For the plates, we always pooled together equal volumes from each well and sampled the same number of plates from each set of 24, so that the pools have equal volumes of culture from each of the 24 populations. For these two pools (plate and flask), we used a Coulter Counter Z2 (Beckman Coulter) to measure the average cell density and pelleted cells for sequencing. After 24 hours, we mixed 16 wells from each population (we achieved this by pooling by column from 2 of the 24 plates per set) and diluted 1:$2^{10}$ into 32 wells (4 new plates per set). We repeated this transfer on 4 plates (2 from each set) left to grow for 29 hours instead of 24, and for both we pooled and pelleted samples after 7 hours of new growth (at 31 and 36 hours).

We processed the sequencing data using the same procedure as for E1 and E2. We estimated cell density for each population at each timepoint using the average cell density from the Coulter Counter and the percentage of reads mapping to each transformation library (fig. S10A), and estimated exponential growth rates for each segregant ($gr_{\mathrm{seg}}$) as the slope of $log_2(cell\ density)$ with respect to hours between 4 and 13 hours.

To calculate fitness effects of mutations, we pooled barcode counts by mutation, only using barcodes that were seen in the corresponding E2 assay and were not excluded as outliers. We do not estimate fitness effects for individual barcodes here because the destructive sampling adds too much noise at the barcode level. We use cell density estimates to determine how many generations have elapsed between two timepoints: $gens_{t1\_t2} = log_2(cell\ density_{t2}) - log_2(cell\ density_{t1})$. We estimated the fitness effect of each mutation using the same log-slope-based method as in E1 and E2, but measured fitness effects over shorter time intervals. We measure $s_{exponential}$ between hours 4-13 (this choice of timing is based on our measured growth curves, see fig. S10A). We measure $s_{lag}$ between hours 0-7 and 24-31 (i.e. the first seven hours of both dilution cycles; we found this provides better estimates of lag phase growth than using shorter periods), and $s_{saturation}$ between hours 20-29 (based on measured growth curves in fig. S10A). We scale the exponential and lag measurements by $gens_{t1\_t2}$ for each time interval, but for the saturation measurement, since cell density changes very little, we simply scale by hours. For the 96-well plate environment, we use an average across replicates for each of these three *s* measures. Because there is growth during the 0-7 hour timeframe, we calculate the "lag effect" per generation by (1) subtracting the measured exponential *s* from the measured lag *s*, (2) multiplying this value by the number of generations in the first 7 hours (determined from cell count data) to get a per-cycle lag effect, and (3) dividing by 10 to get a scaled per-generation lag effect that is comparable to the *s* values measured in E1 and E2:

$$lag\ effect = \frac{(s_{lag} - s_{exponential}) * gens_{0hr\_7hr}}{10}.$$

Similarly, to calculate the "saturation effect," we (1) multiply the saturation *s* by the number of hours spent in saturation (determined from cell count data) and (2) divide by 10 to get a scaled per-generation effect:

$$saturation\ effect = \frac{s_{saturation} * hours_{saturation}}{10}$$, where $hours_{saturation}$ is estimated using the growth rates estimated from cell density data, assuming 10 generations of growth to reach saturation and no lag: $hours_{saturation} = 24 - 10/gr_{\text{seg}}$.

**Growth curve experiment results**

The exponential growth rates of segregants we measured using cell counts from the growth curve (GC) experiment in plates or in flasks are strongly correlated with competitive fitness measured during batch culture (fig. S10B). We find that our measures of the fitness effects of mutations during exponential phase ("exponential *s*") correlate strongly with the fitness effects we measured in E2 (fig. S11B). Including the lag effect improves the correlation slightly, but including a saturation effect does not (fig. S11B). Using the residual measurement (E2 *s* - exponential *s*), we can ask whether any segregants or mutations have more non-exponential fitness effects. The mean residual for each segregant goes up slightly with background fitness, but the positive slope indicates that the pattern of worse fitness effects in more fit strains would, if anything, be stronger if we only considered exponential growth (fig. S11A). We next asked if any mutations had residuals across segregants that deviated significantly from zero using one sample t-tests and a Benjamini-Hochberg multiple hypothesis correction. We find one mutation that has systematically higher fitness effects during exponential phase in our GC experiment, in MPC2, and a deleterious effect in lag or saturation phase explains this deviation (fig. S11C). Excluding the MPC2 mutation from our analysis does not change any of our conclusions. Most importantly, if we plot fitness effects measured during exponential phase against background fitness (as in Fig. 3 in the main text), we see that the patterns of background fitness or QTL dependence found in the E2 experiment are preserved (fig. S12). This is the case even for fitness effects measured during exponential phase of growth in the shaken flask, plotted against the exponential growth rate measured from the cell density data for the flask experiment (fig. S12).

#### Data and Code Availability

Raw sequencing data and processed count data will be submitted to GenBank and Dryad, respectively, before publication. All analysis code is available at https://github.com/mjohnson11/TnSeq_Pipeline .

### Figures

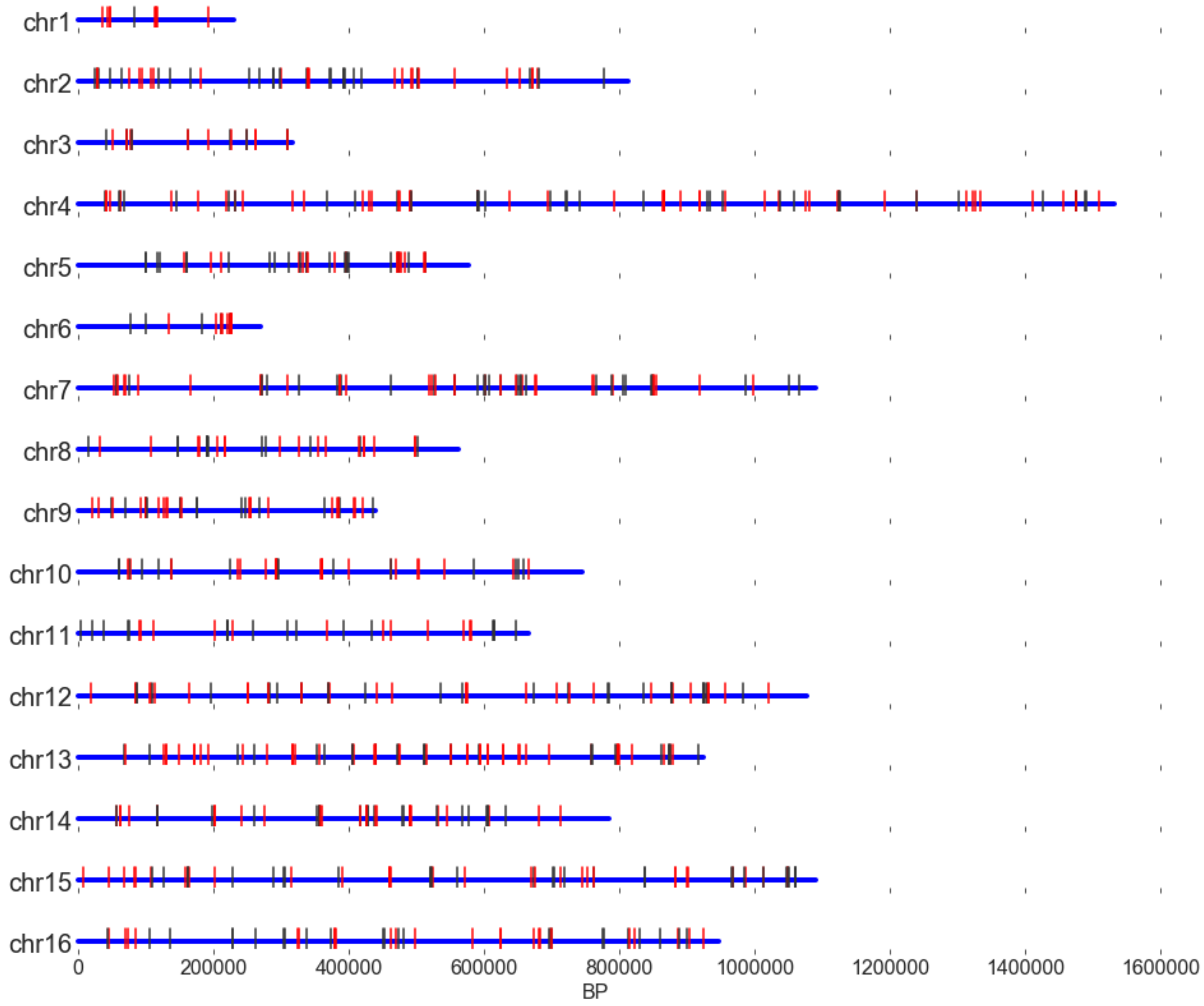

**Fig. S1.** Map of Tn7 insertions with measured fitness effects in E1. Each blue line represents a chromosome. Red lines indicate insertions in coding regions, black lines represent insertions in intergenic regions.

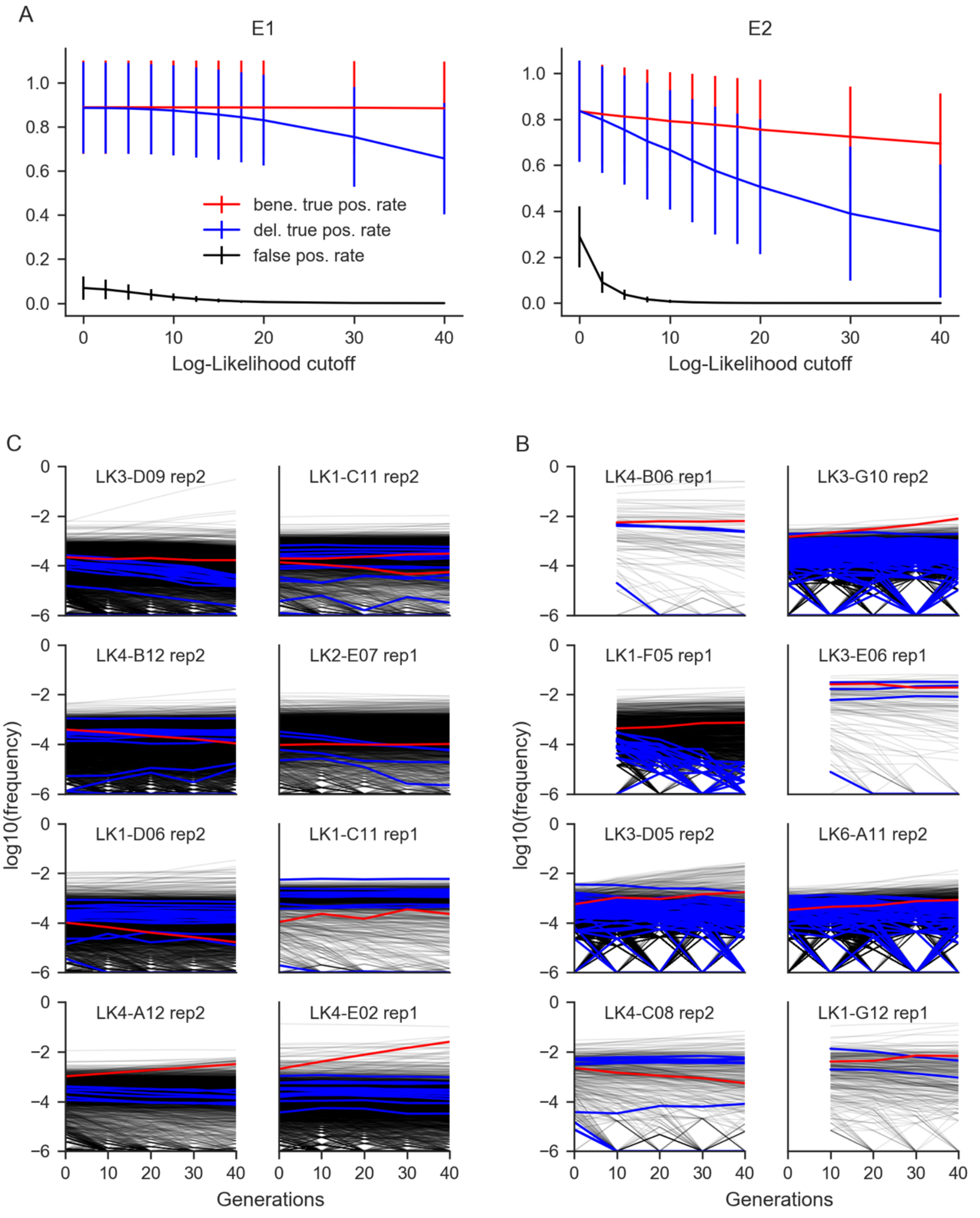

**Fig. S2. A.** True positive (outlier correctly excluded) and false positive (correct barcode excluded) rates (Y axis) for different log-likelihood ratio cutoffs (X axis) for simulated data based on E1 (left) and E2 (right). **B.** Examples from E1 of excluded outlier barcode frequency trajectories (red) compared with other barcode trajectories for the same mutation (blue). The x axis is generations and the y-axis is log10(frequency). **C.** Same as (B) for E2.

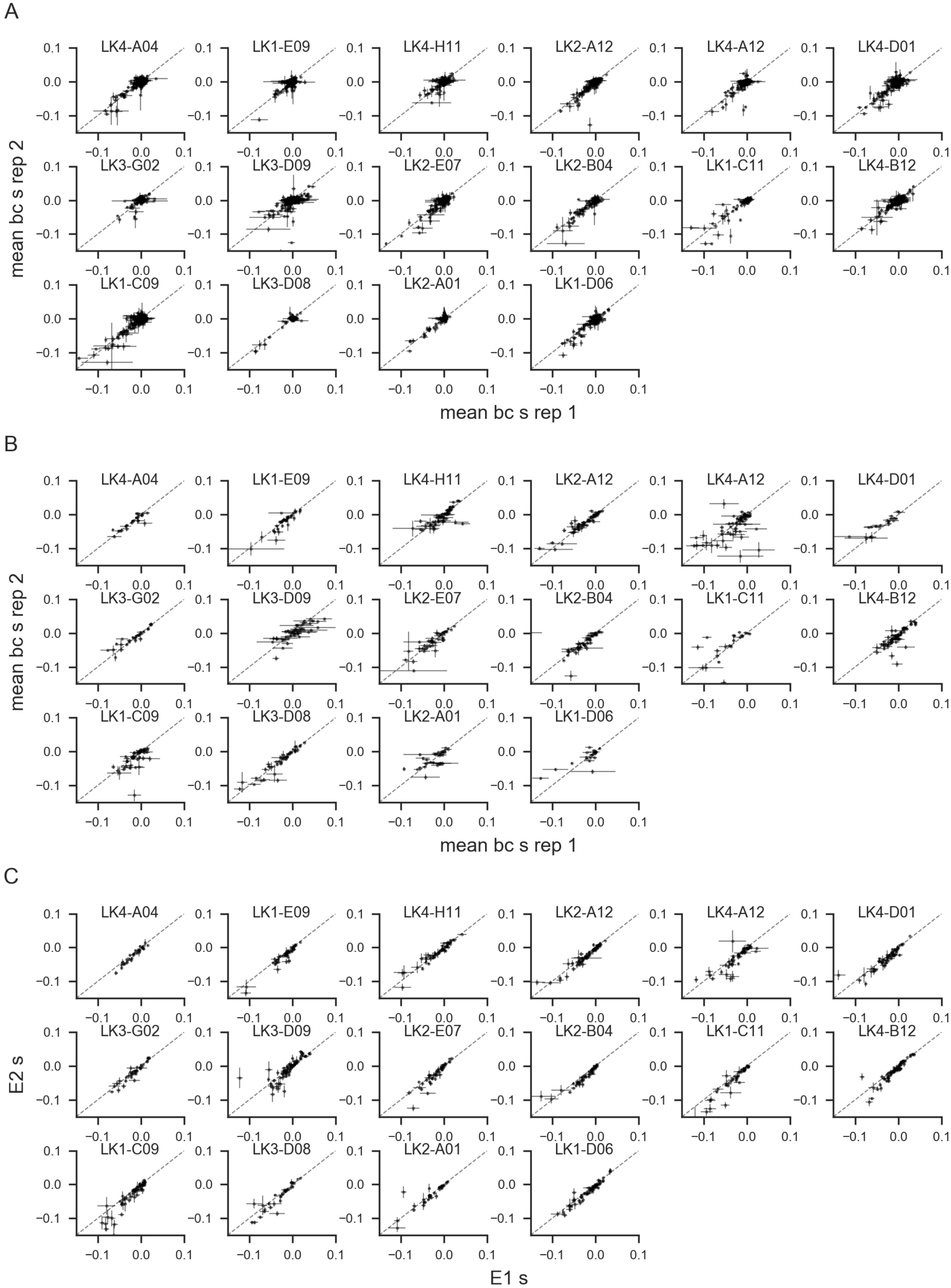
A
LK4-A04
LK1-E09
LK4-H11
LK2-A12
LK4-A12
LK4-D01
LK3-G02
LK3-D09
LK2-E07
LK2-B04
LK1-C11
LK4-B12
LK1-C09
LK3-D08
LK2-A01
LK1-D06
mean bc s rep 2
mean bc s rep 1
B
LK4-A04
LK1-E09
LK4-H11
LK2-A12
LK4-A12
LK4-D01
LK3-G02
LK3-D09
LK2-E07
LK2-B04
LK1-C11
LK4-B12
LK1-C09
LK3-D08
LK2-A01
LK1-D06
mean bc s rep 2
mean bc s rep 1
C
LK4-A04
LK1-E09
LK4-H11
LK2-A12
LK4-A12
LK4-D01
LK3-G02
LK3-D09
LK2-E07
LK2-B04
LK1-C11
LK4-B12
LK1-C09
LK3-D08
LK2-A01
LK1-D06
E2 s
E1 s

**Fig. S3**. Correlations between fitness effect measurements for the 16 segregants for which we have sufficient fitness effect data in both E1 and E2. Each panel corresponds to a single segregant, and error bars are standard errors. **A.** Correlations between replicate s measurements for E1 for all mutations with at least 2 barcodes in each replicate. **B.** Correlations between replicate s measurements for E2 for all mutations with at least 2 barcodes in each replicate. **C.** Correlations between s measurements in E1 and E2 for all mutations with at least 3 barcodes in each experiment.

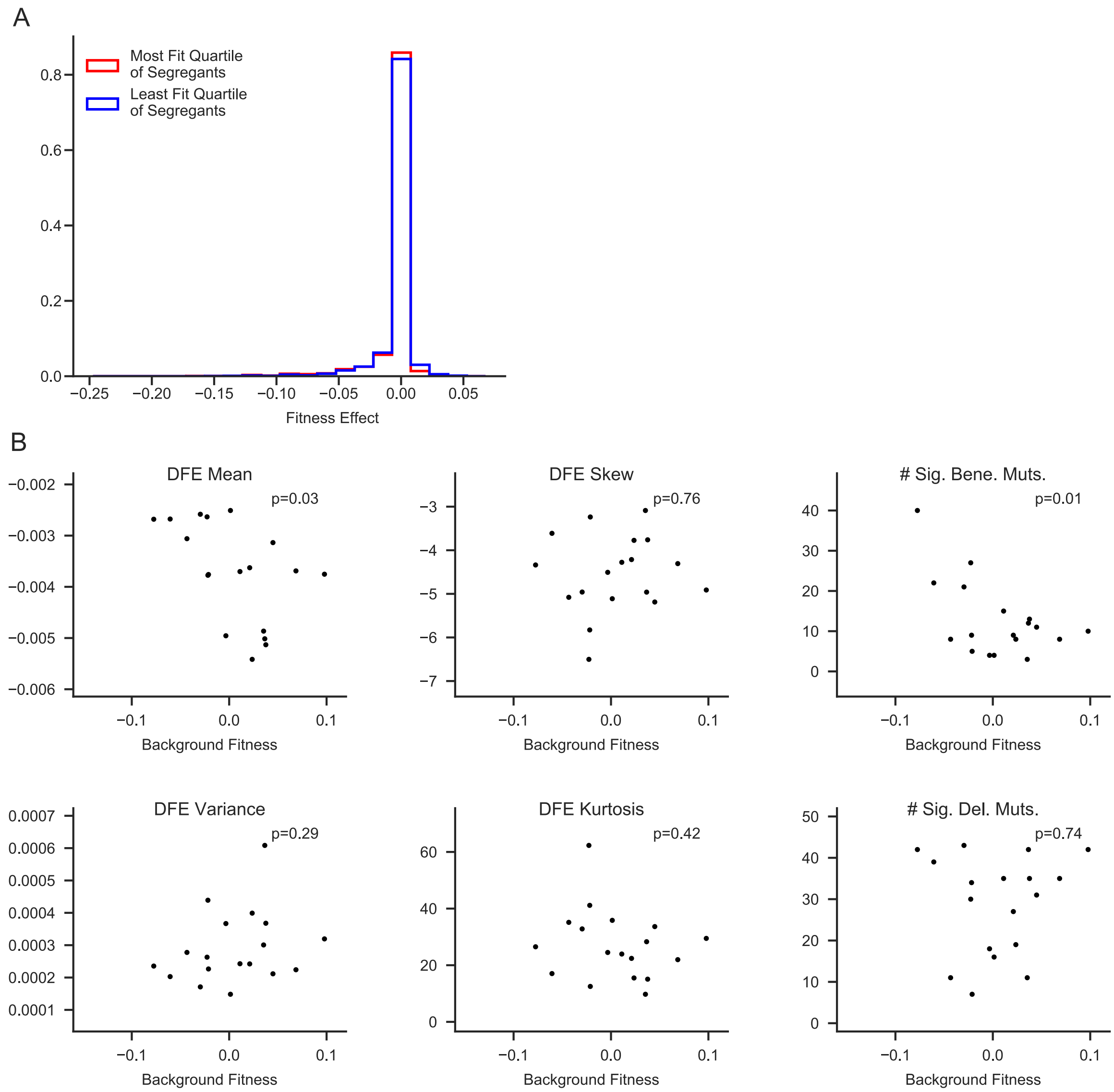

**Fig. S4. A.** Combined distribution of fitness effects of the most-fit and least-fit quartiles of segregants in E1. **B.** DFE statistics as a function of background fitness for E1. P-values for linear regressions against background fitness are shown. The methods used to generate these statistics are described in the "Modeling genetic determinants of DFE statistics" section.

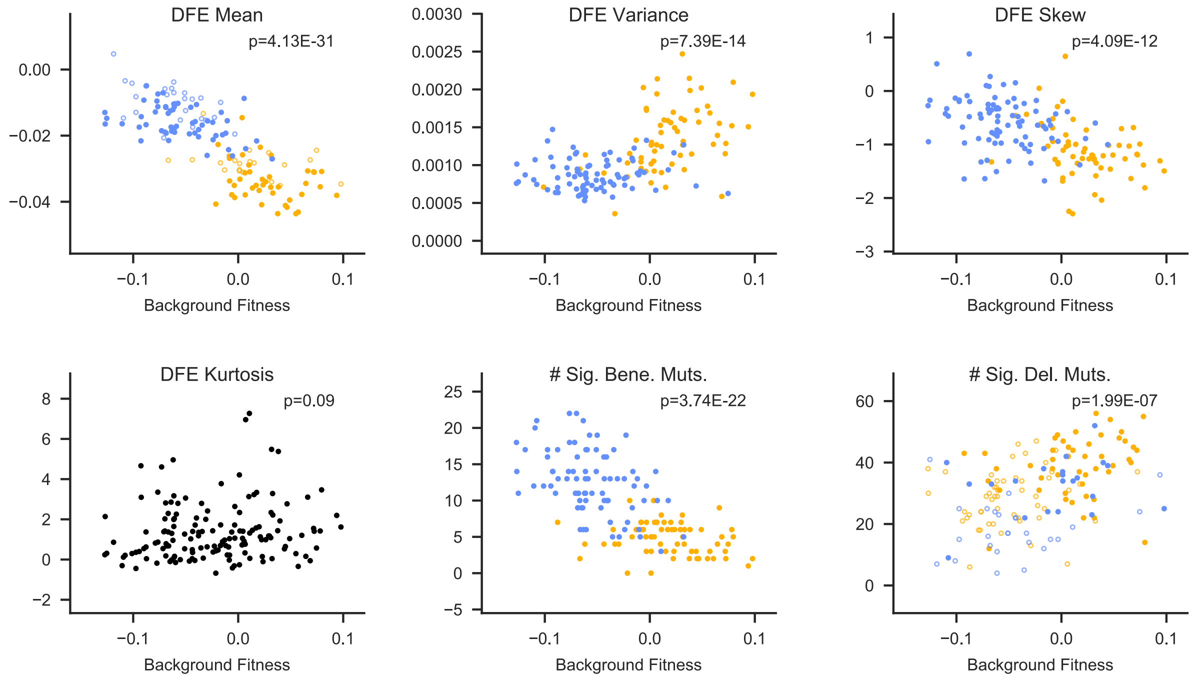

**Fig. S5.** DFE statistics as a function of background fitness for E2. P-values for linear regressions against background fitness are shown. Allelic state at the largest-effect quantitative trait locus for the DFE statistic is shown by yellow (BY) or blue (RM) color; allelic state at the second largest-effect quantitative trait locus is shown by closed (BY) or open (RM) symbol. The methods used to generate these statistics are described in the "Modeling genetic determinants of DFE statistics" section.

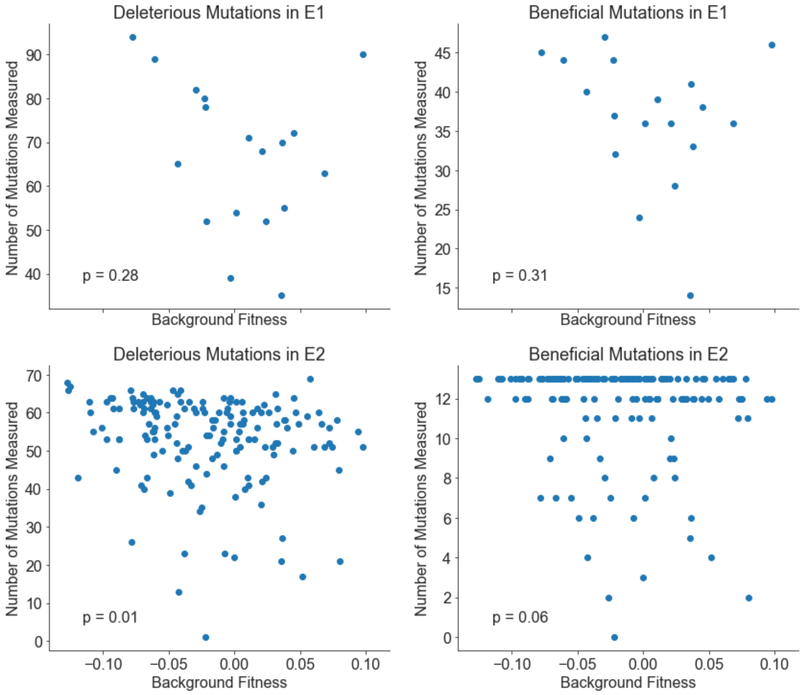

**Fig. S6.** The number of deleterious and beneficial mutations measured in E1 (top) do not vary significantly as a function of background fitness. In E2 (bottom), the number of deleterious mutations measured decreases slightly with increasing background fitness. This suggests that the measurements we are missing would only strengthen our claims about shifting DFEs (see "Missing fitness measurements" for details).

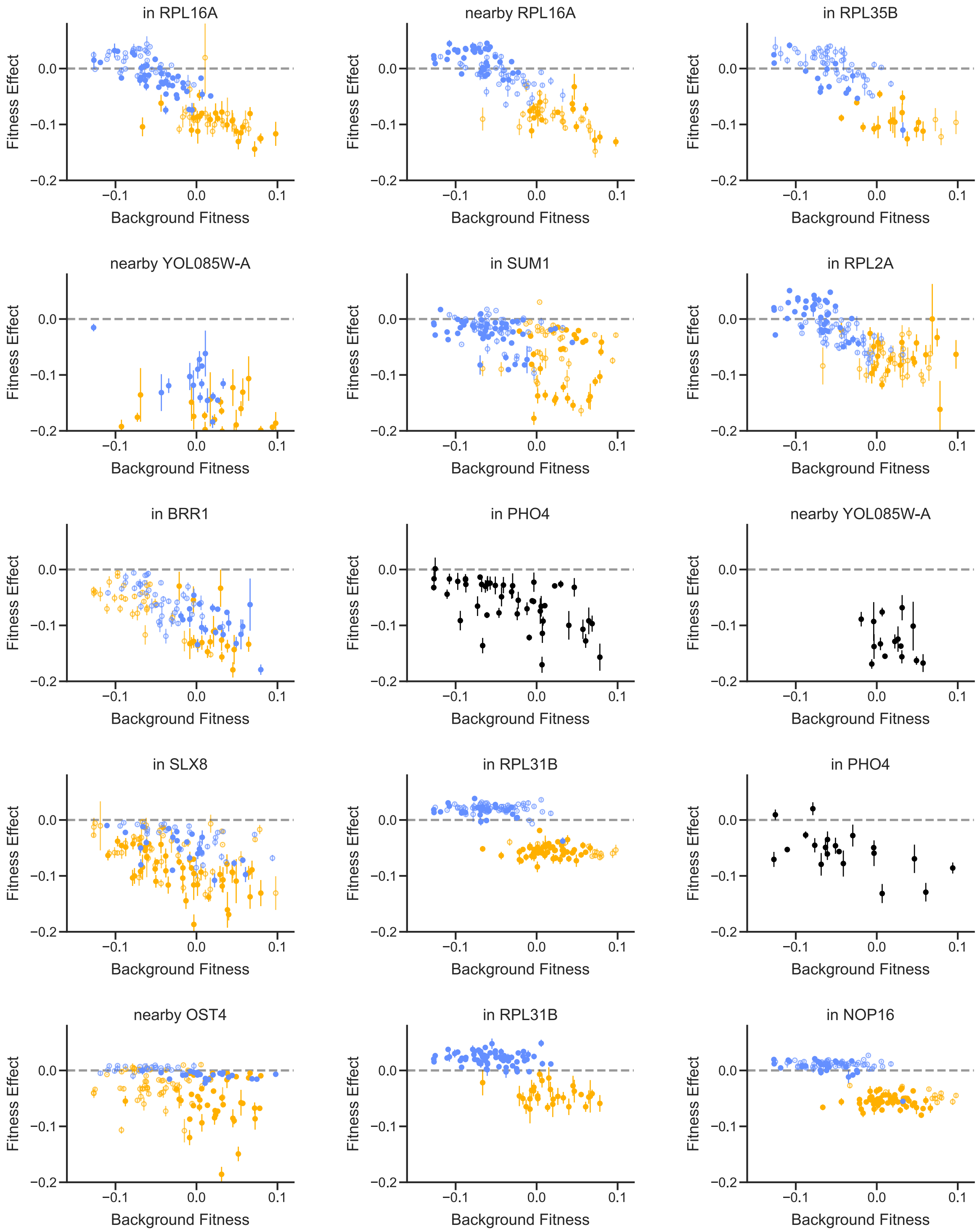
in RPL16A
nearby RPL16A
in RPL35B
nearby YOL085W-A
in SUM1
in RPL2A
in BRR1
in PHO4
nearby YOL085W-A
in SLX8
in RPL31B
in PHO4
nearby OST4
in RPL31B
in NOP16
Fitness Effect
Background Fitness
0.0
−0.1
−0.2
0.1

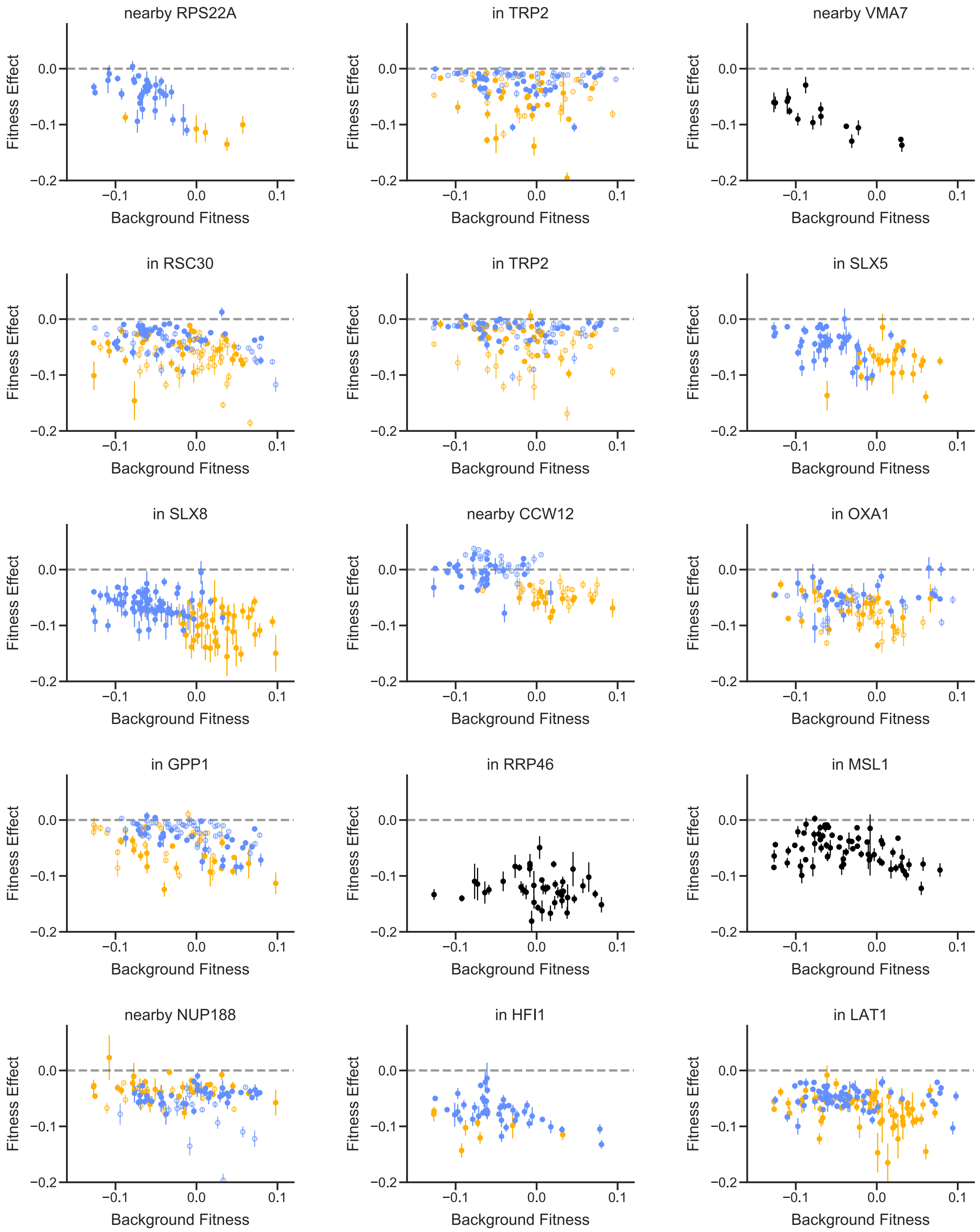
nearby RPS22A
in TRP2
nearby VMA7
in RSC30
in TRP2
in SLX5
in SLX8
nearby CCW12
in OXA1
in GPP1
in RRP46
in MSL1
nearby NUP188
in HFI1
in LAT1
Fitness Effect
Background Fitness
0.0
−0.1
−0.2
−0.1
0.0
0.1

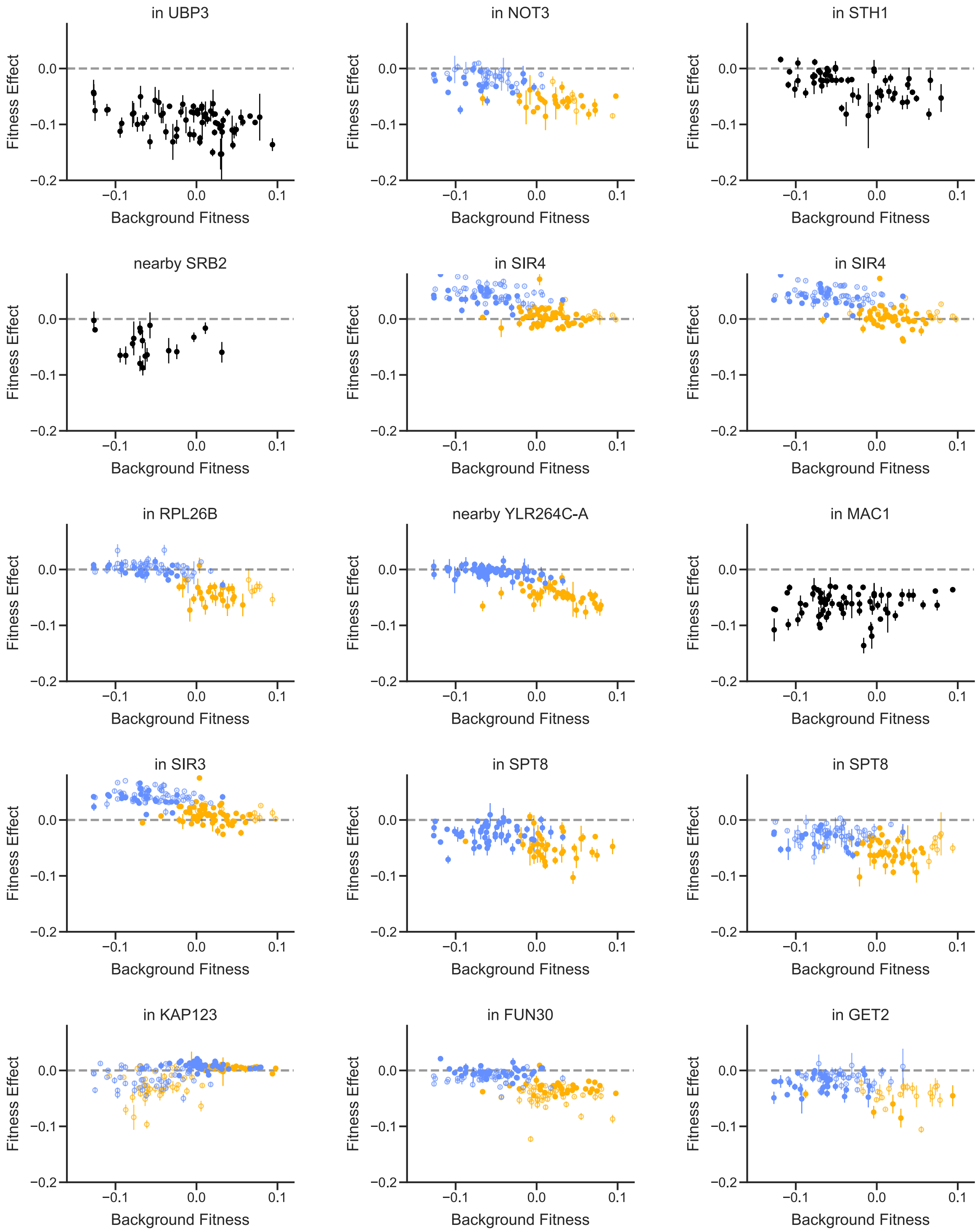
in UBP3
in NOT3
in STH1
nearby SRB2
in SIR4
in SIR4
in RPL26B
nearby YLR264C-A
in MAC1
in SIR3
in SPT8
in SPT8
in KAP123
in FUN30
in GET2
Fitness Effect
Background Fitness
0.0
−0.1
−0.2
−0.1
0.0
0.1

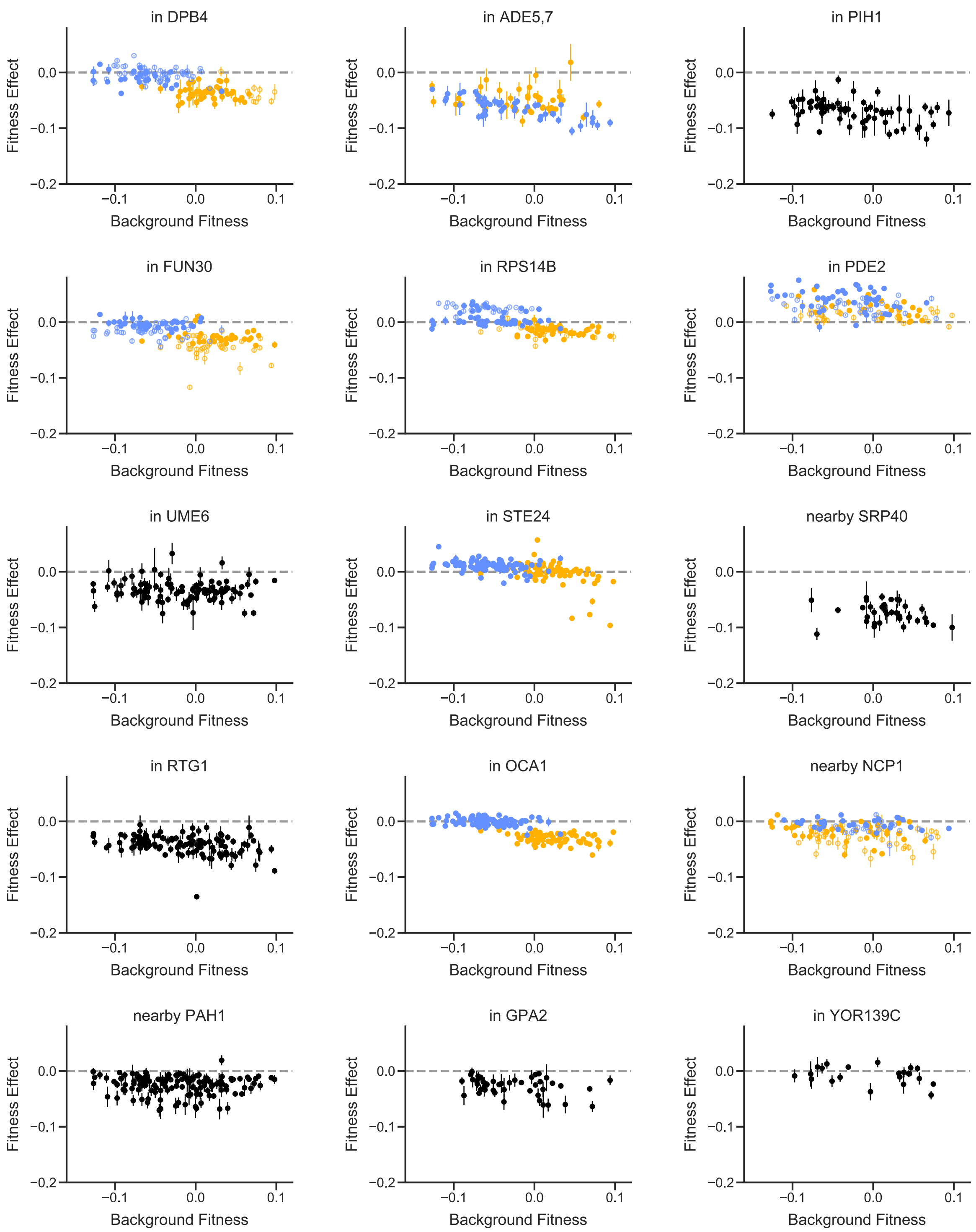
in DPB4
in ADE5,7
in PIH1
in FUN30
in RPS14B
in PDE2
in UME6
in STE24
nearby SRP40
in RTG1
in OCA1
nearby NCP1
nearby PAH1
in GPA2
in YOR139C
Fitness Effect
0.0
−0.1
−0.2
Background Fitness
−0.1
0.0
0.1

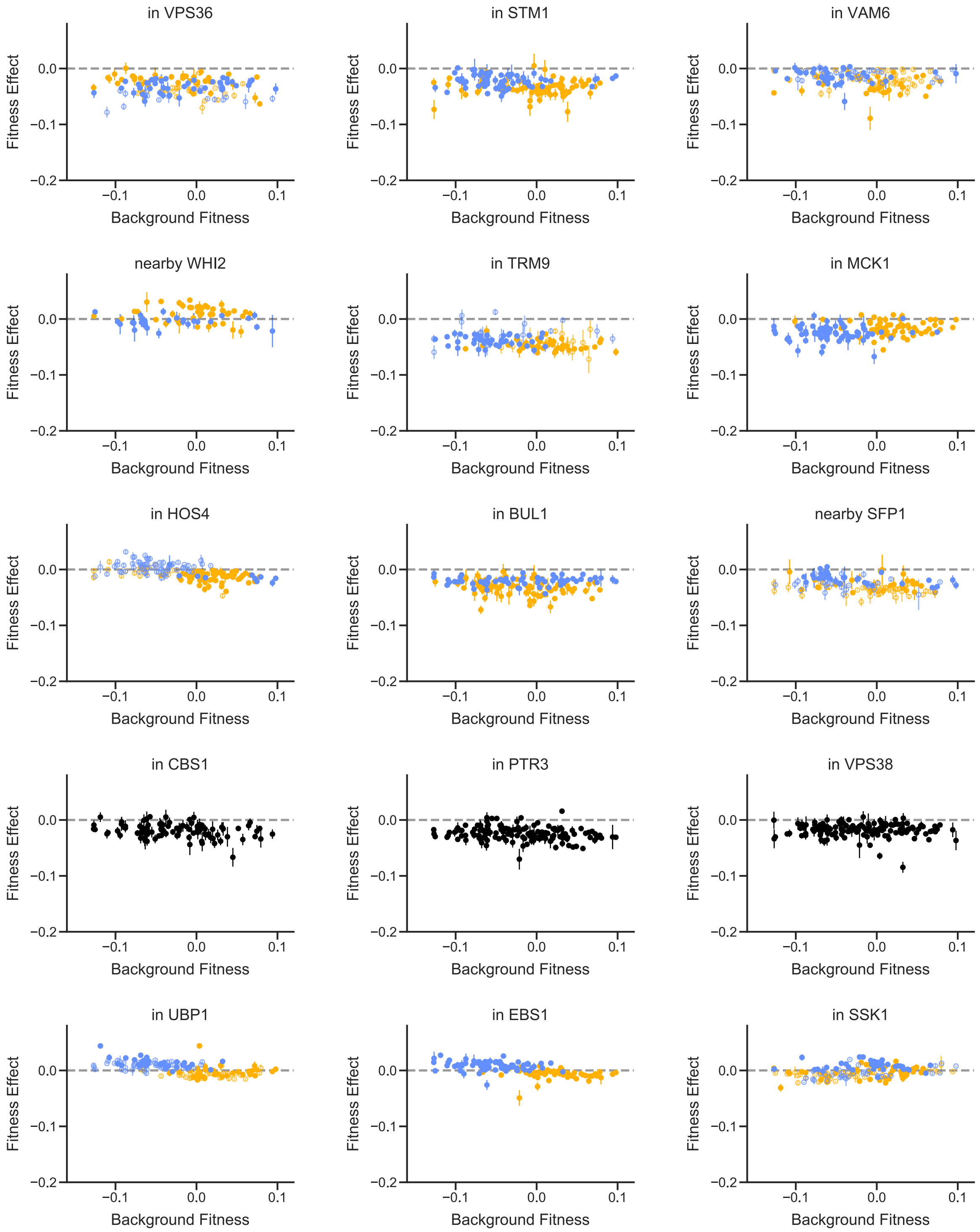
in VPS36
in STM1
in VAM6
nearby WHI2
in TRM9
in MCK1
in HOS4
in BUL1
nearby SFP1
in CBS1
in PTR3
in VPS38
in UBP1
in EBS1
in SSK1
Fitness Effect
Background Fitness
0.0
−0.1
−0.2
0.1

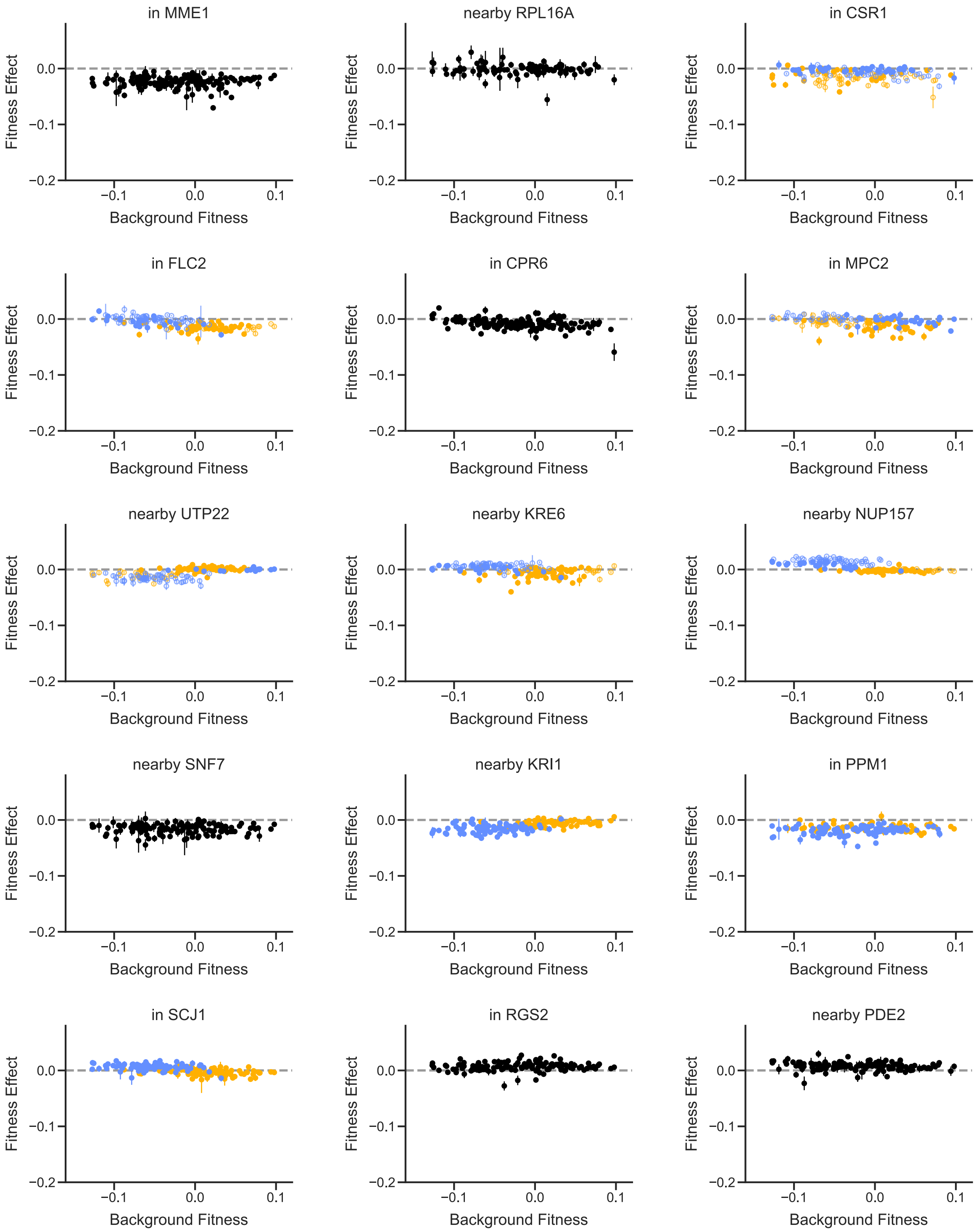
in MME1
nearby RPL16A
in CSR1
in FLC2
in CPR6
in MPC2
nearby UTP22
nearby KRE6
nearby NUP157
nearby SNF7
nearby KRI1
in PPM1
in SCJ1
in RGS2
nearby PDE2
Fitness Effect
Background Fitness
0.0
−0.1
−0.2
0.1

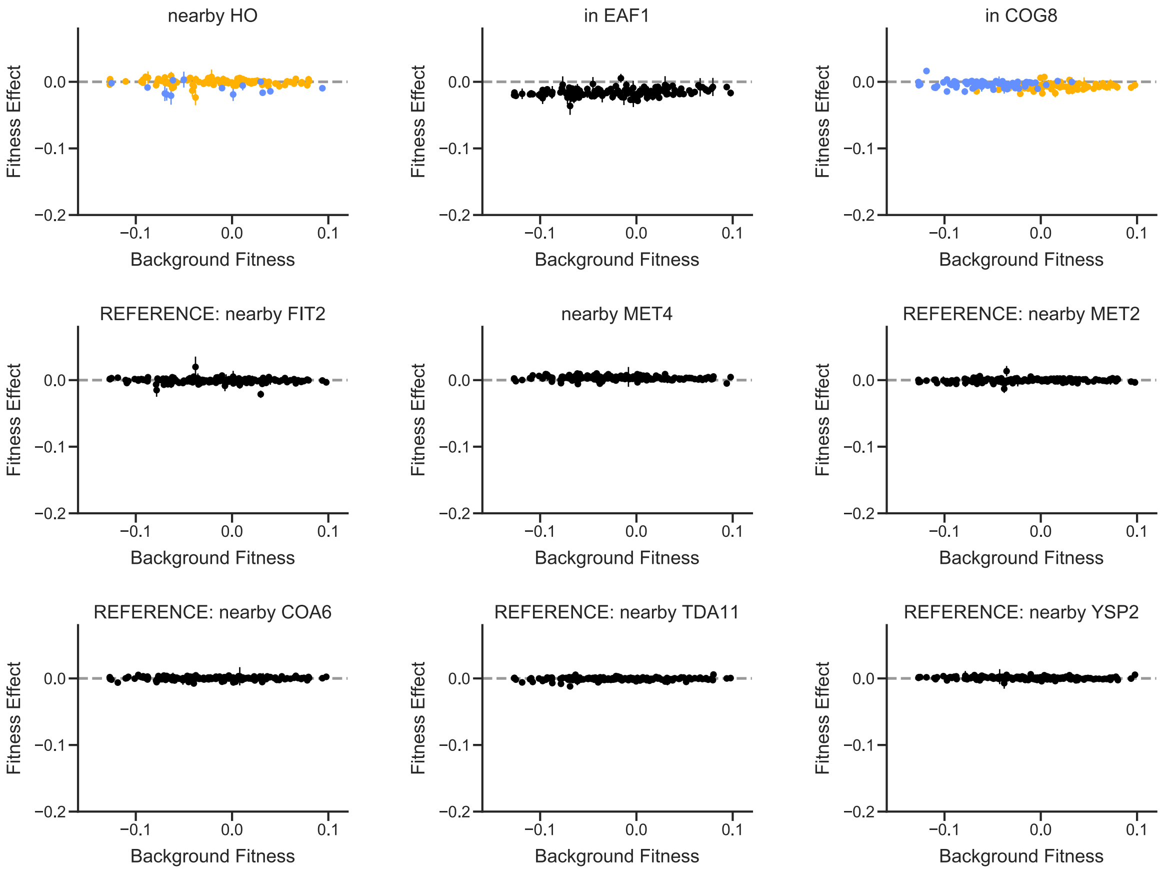

**Fig. S7.** Patterns of epistasis for individual mutations. Fitness effects of all insertion mutations in E2 are plotted against segregant background fitness. Allelic state at the largest-effect quantitative trait locus for the fitness effect of each mutation is shown by yellow (BY) or blue (RM) color; allelic state at the second largest-effect quantitative trait locus is shown by closed (BY) or open (RM) symbol. Error bars represent standard errors (Methods).

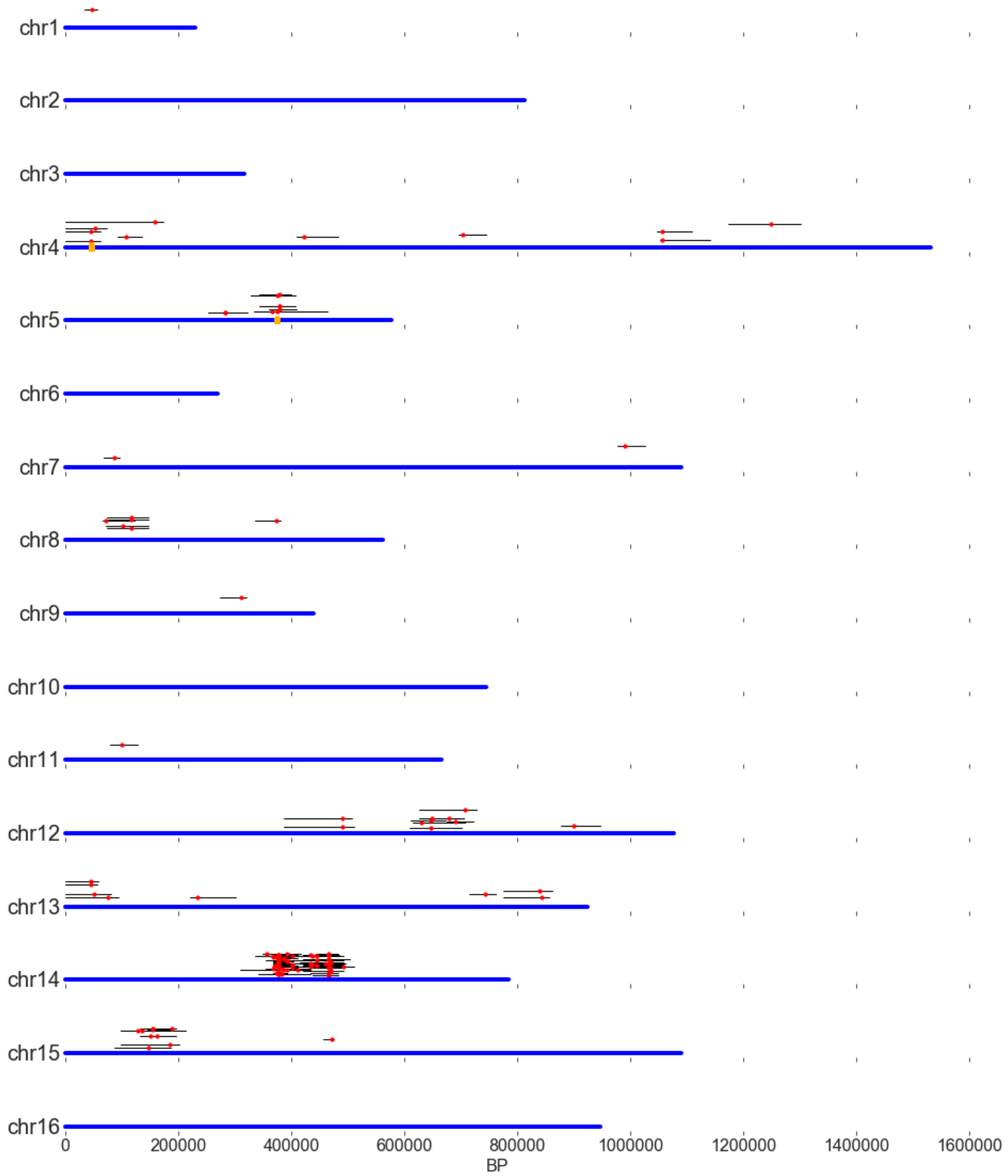

**Fig. S8.** Map of detected QTLs and confidence intervals (red dots and black lines). Each blue line represents a chromosome. Points are randomly displaced vertically to distinguish overlapping QTLs. Yellow blocks represent the antibiotic resistance markers at HO (chr4) and FLO8 (chr5).

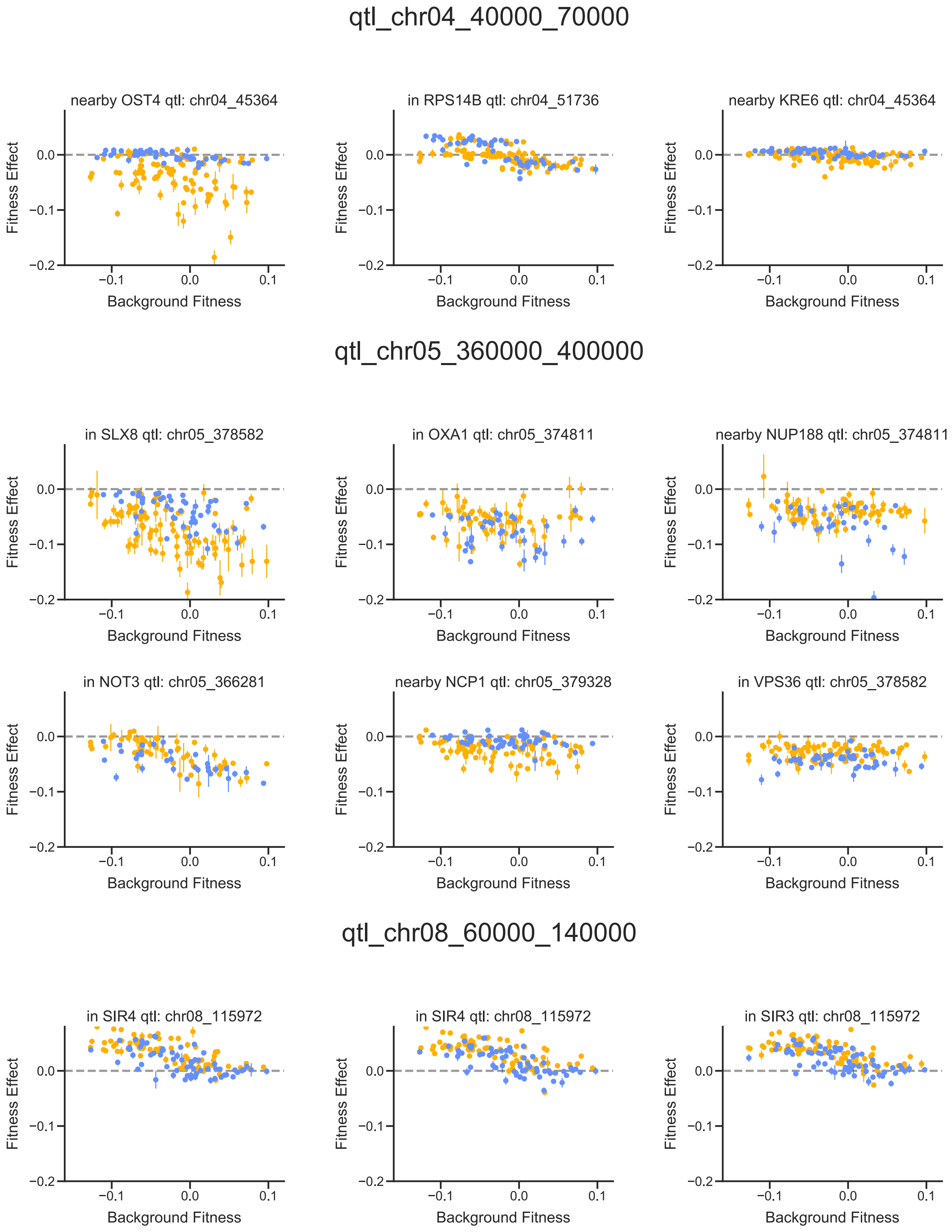
qtl_chr04_40000_70000
nearby OST4 qtl: chr04_45364
in RPS14B qtl: chr04_51736
nearby KRE6 qtl: chr04_45364
qtl_chr05_360000_400000
in SLX8 qtl: chr05_378582
in OXA1 qtl: chr05_374811
nearby NUP188 qtl: chr05_374811
in NOT3 qtl: chr05_366281
nearby NCP1 qtl: chr05_379328
in VPS36 qtl: chr05_378582
qtl_chr08_60000_140000
in SIR4 qtl: chr08_115972
in SIR4 qtl: chr08_115972
in SIR3 qtl: chr08_115972
Fitness Effect
Background Fitness
0.0
−0.1
−0.2
0.1

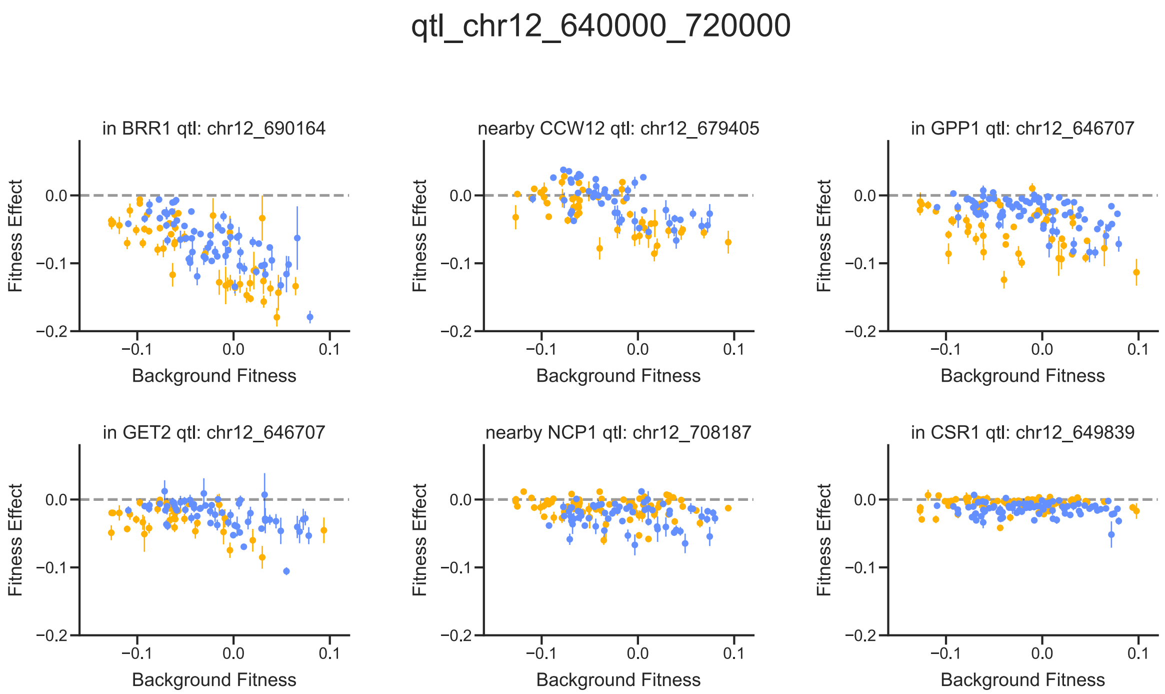
qtl_chr12_640000_720000
in BRR1 qtl: chr12_690164
nearby CCW12 qtl: chr12_679405
in GPP1 qtl: chr12_646707
in GET2 qtl: chr12_646707
nearby NCP1 qtl: chr12_708187
in CSR1 qtl: chr12_649839
Fitness Effect
Background Fitness
0.0
−0.1
−0.2
0.1

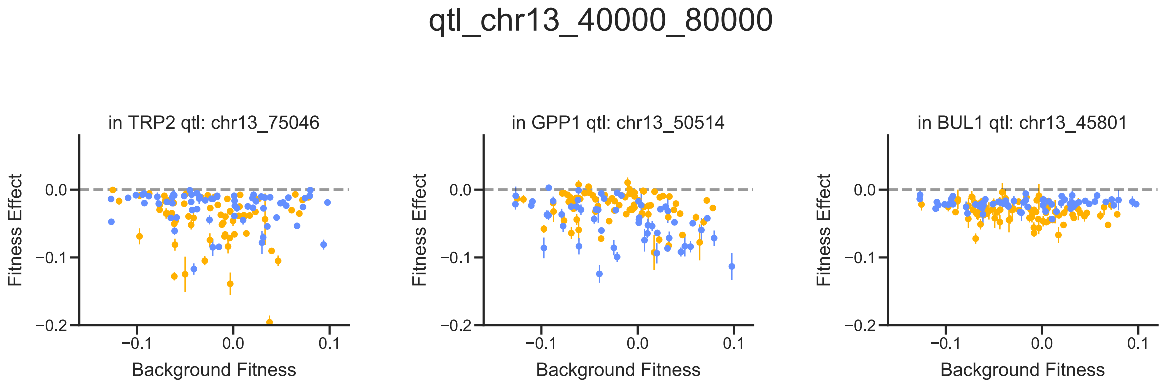
qtl_chr13_40000_80000
in TRP2 qtl: chr13_75046
in GPP1 qtl: chr13_50514
in BUL1 qtl: chr13_45801
Fitness Effect
Background Fitness
0.0
−0.1
−0.2
0.1

qtl_chr14_360000_420000

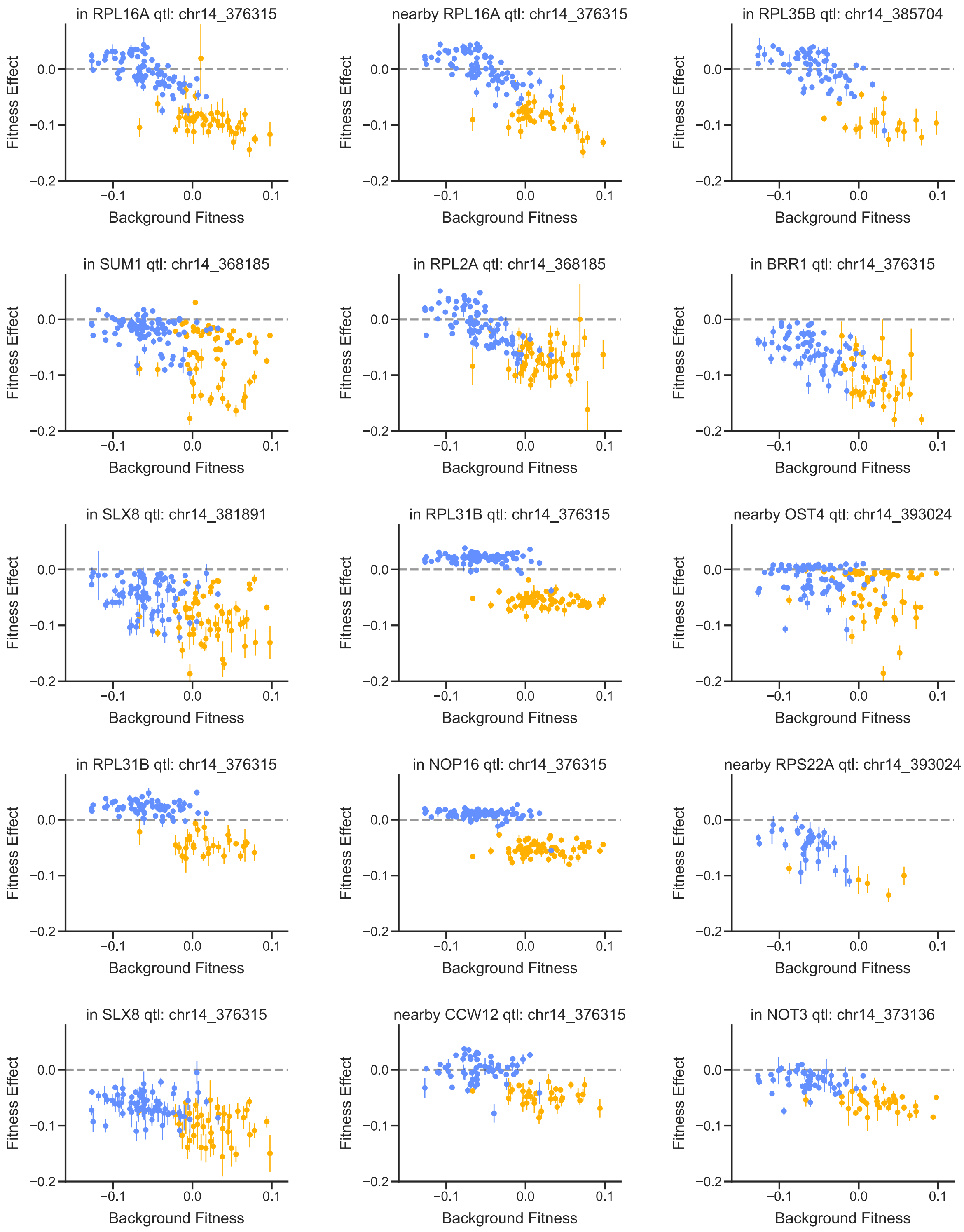

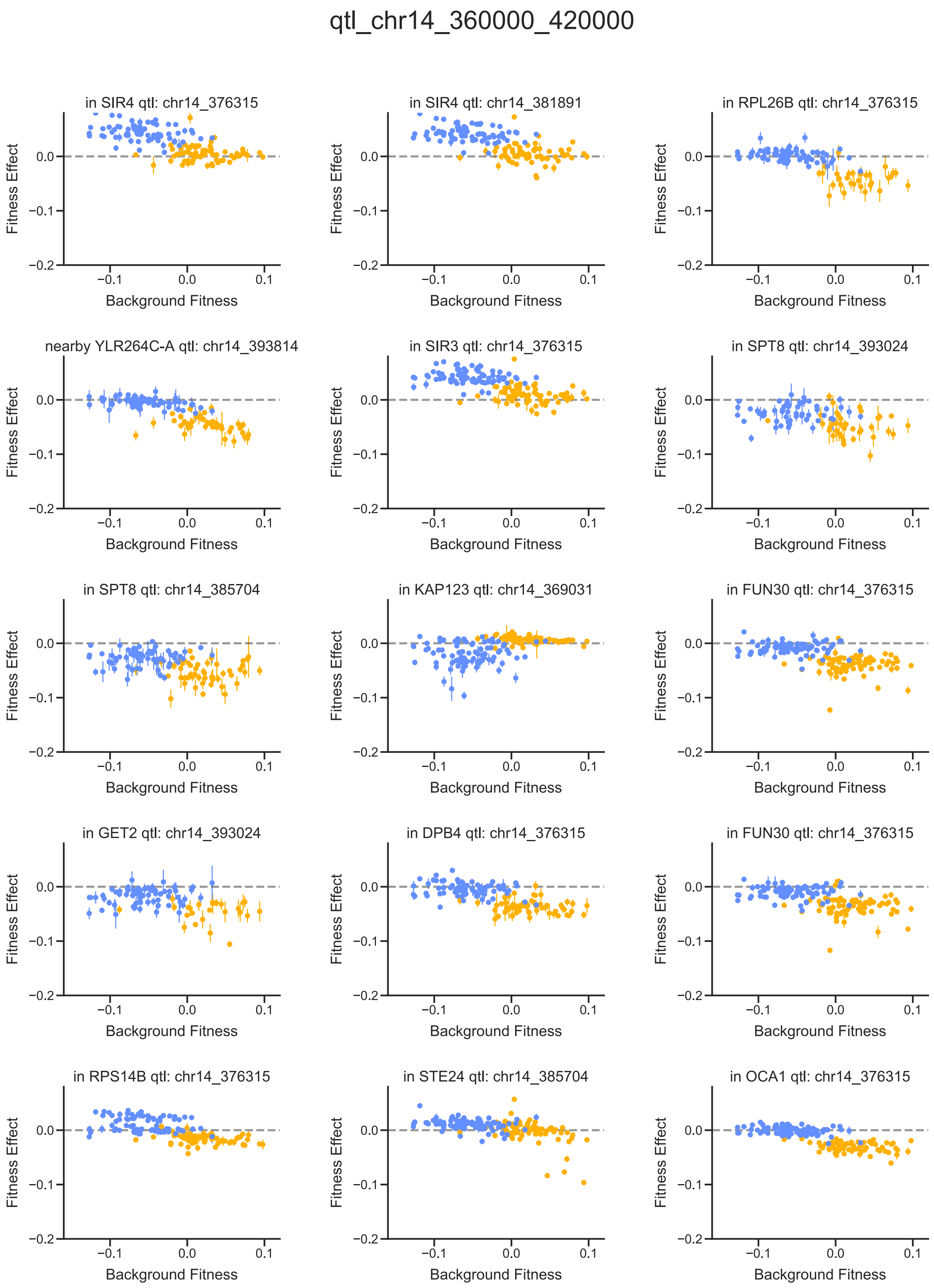
qtl_chr14_360000_420000
in SIR4 qtl: chr14_376315
in SIR4 qtl: chr14_381891
in RPL26B qtl: chr14_376315
nearby YLR264C-A qtl: chr14_393814
in SIR3 qtl: chr14_376315
in SPT8 qtl: chr14_393024
in SPT8 qtl: chr14_385704
in KAP123 qtl: chr14_369031
in FUN30 qtl: chr14_376315
in GET2 qtl: chr14_393024
in DPB4 qtl: chr14_376315
in FUN30 qtl: chr14_376315
in RPS14B qtl: chr14_376315
in STE24 qtl: chr14_385704
in OCA1 qtl: chr14_376315
Fitness Effect
Background Fitness
0.0
−0.1
−0.2
0.1

#### qtl_chr14_360000_420000

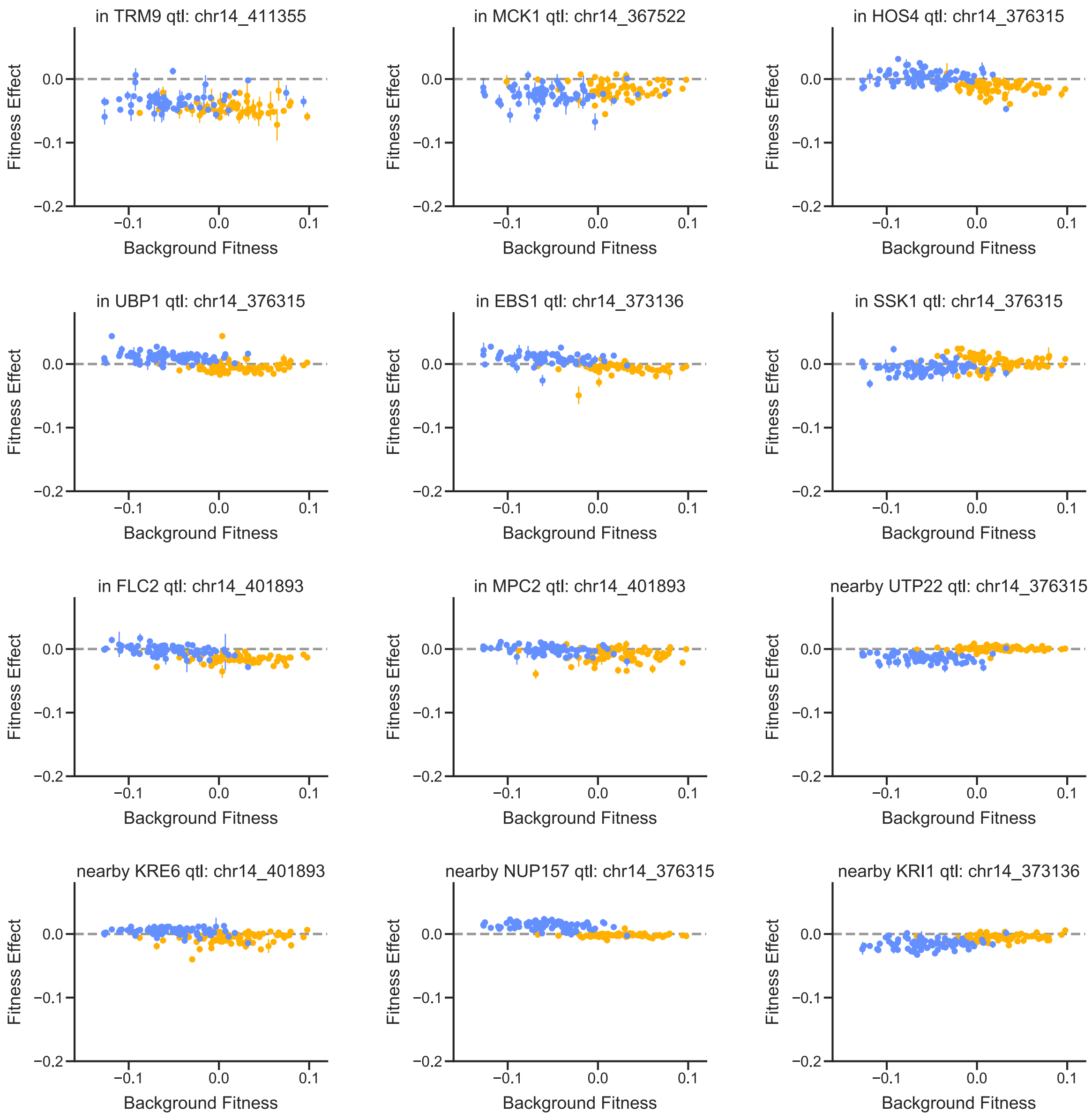

qtl_chr14_430000_500000

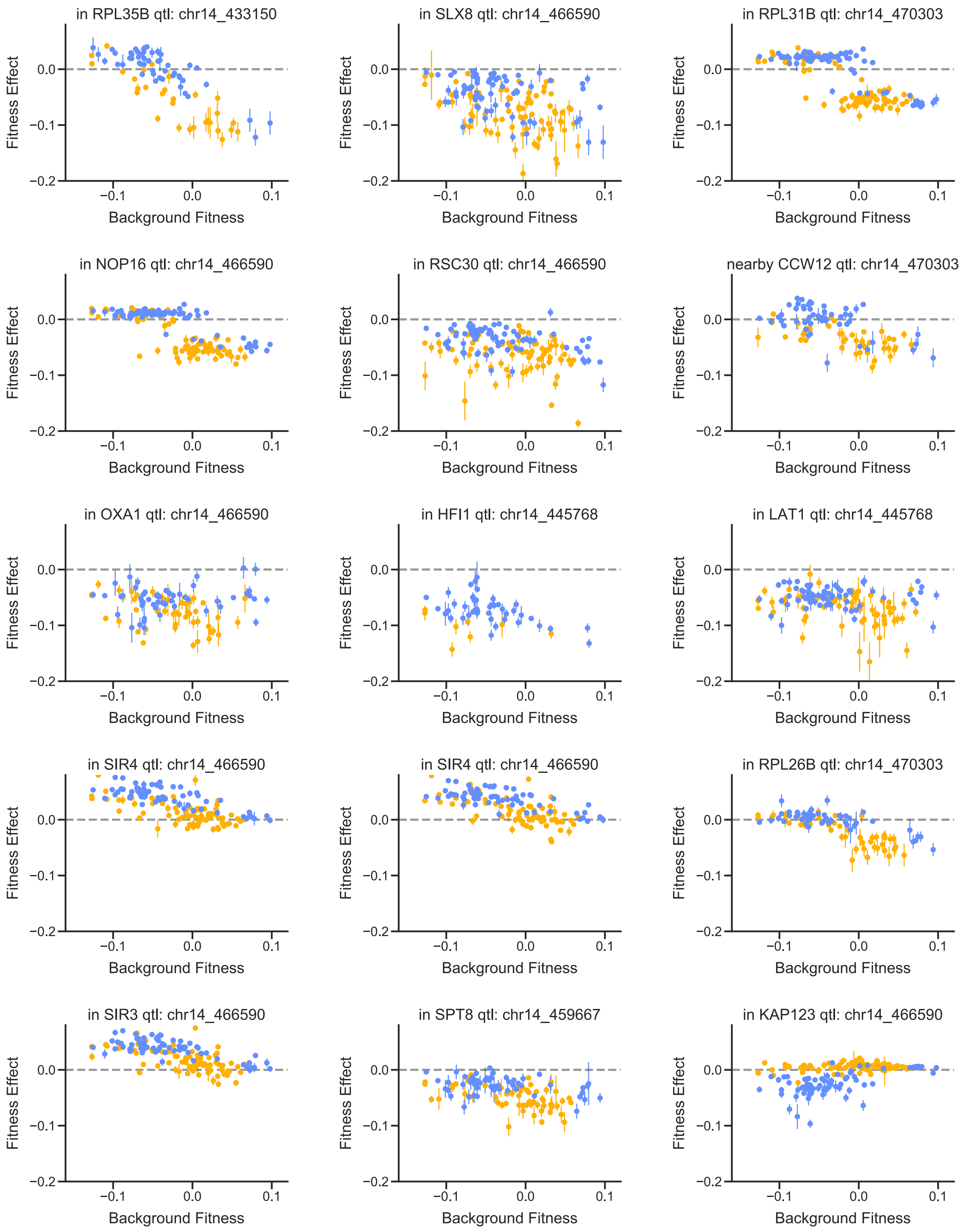

qtl_chr14_430000_500000

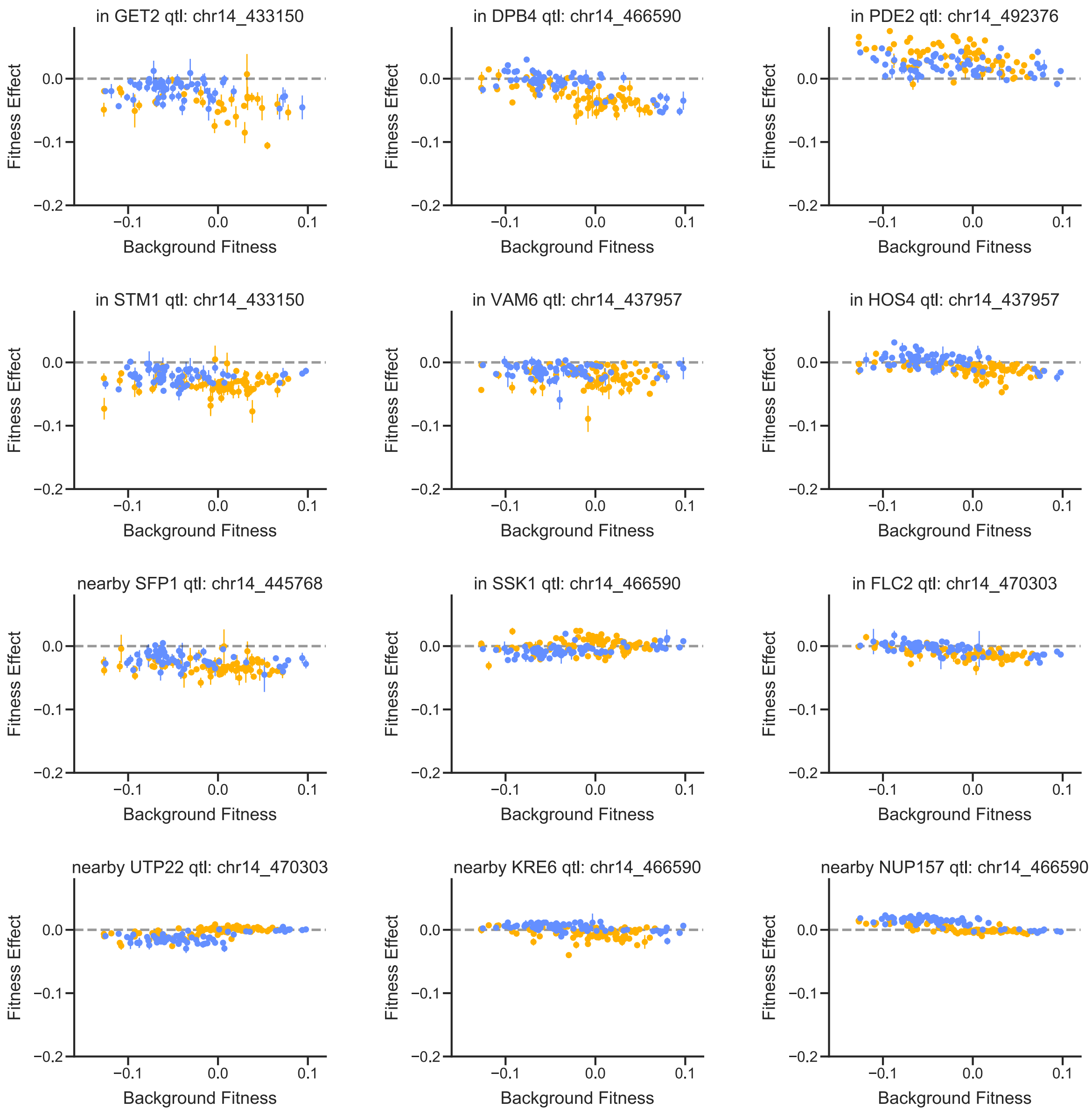

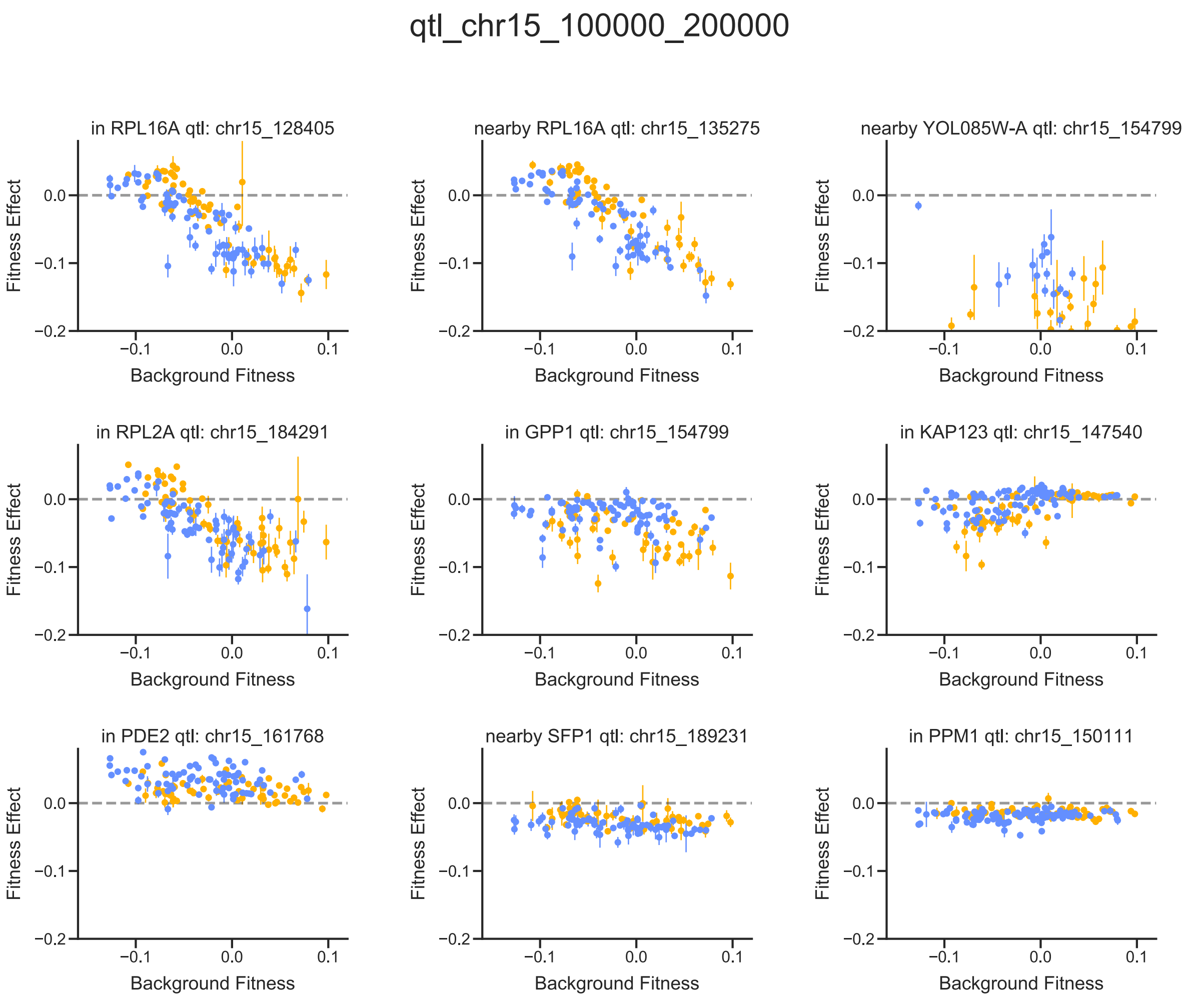

**Fig. S9.** For each multi-hit QTL region, we plot all mutations with fitness effects affected by a QTL in that region. For each plot, the allelic state at the QTL in question is shown by yellow (BY) or blue (RM) color. Note that mutations may appear more than once in this figure if they are affected by multiple QTLs. Also note that the first two multi-hit QTL regions are near the antibiotic resistance cassettes and are discussed in "Multi-hit QTLs and GO-Term enrichment" (chr04 40000-70000 QTL is near hoΔ::hphMX4 and chr05 360000-400000 QTL is near flo8Δ::natMX4). Error bars represent standard errors.

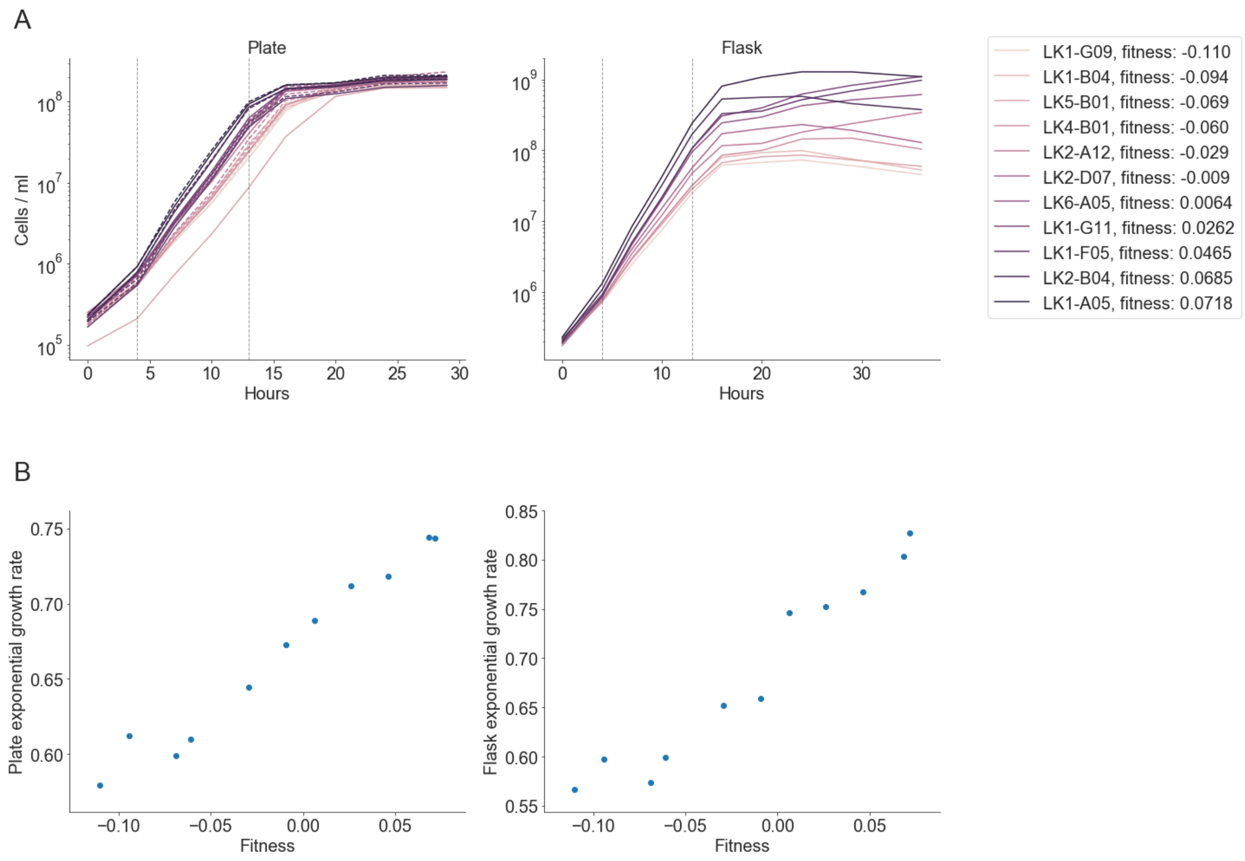

**Fig. S10. A.** Cell density over time in the growth curve experiment, measured using Coulter Counter cell counts and the relative frequency of barcodes corresponding to different populations over time. In the left panel, dotted lines represent replicate 2 for each segregant (only one replicate was included in the flask experiment, right panel). The time period between vertical dotted lines was used to estimate exponential growth rates and was the period used to measure fitness effects during exponential phase. **B.** Exponential growth rate in the plate and flask environment correlates with competitive fitness measured in batch culture.

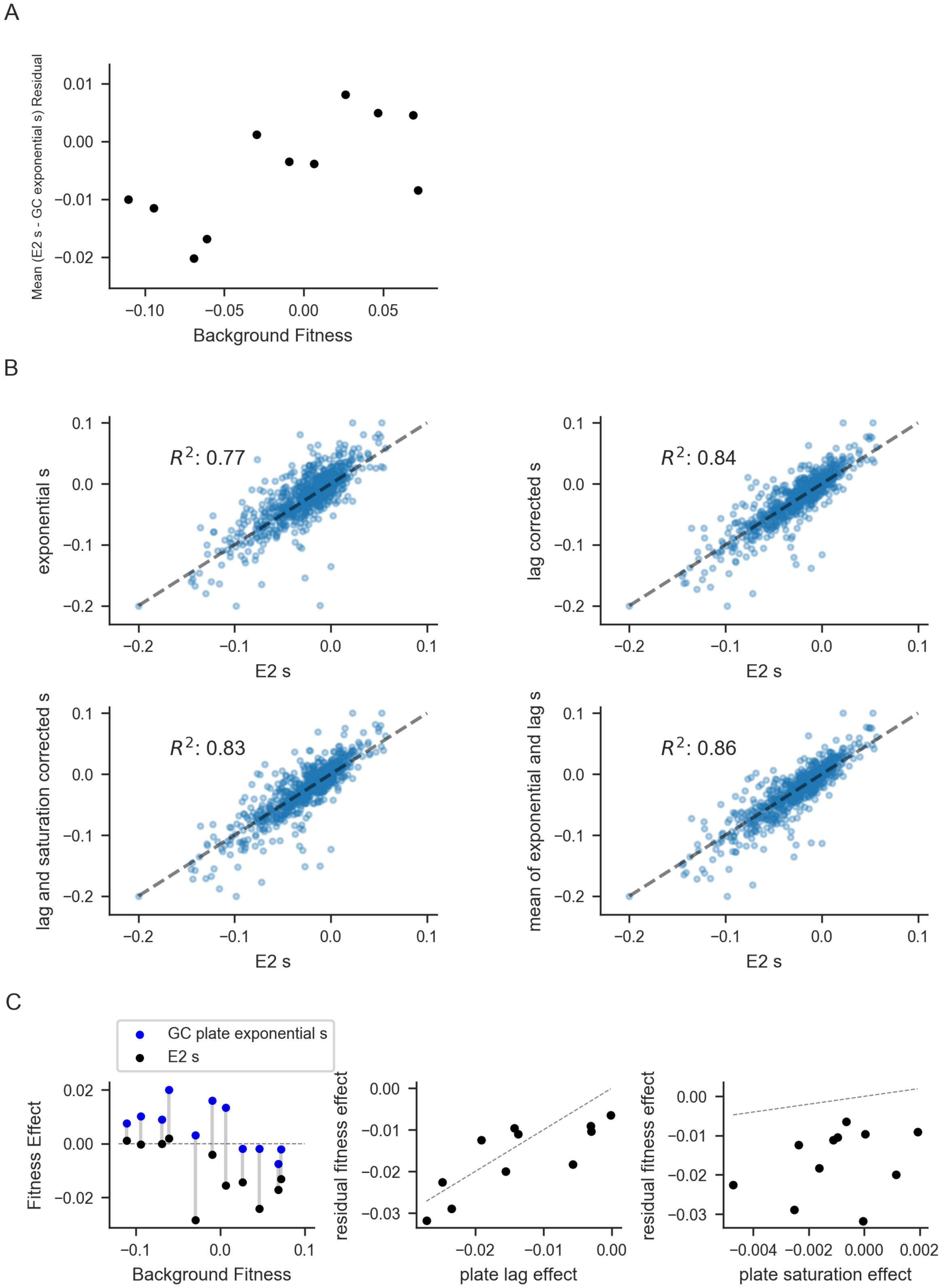

**Fig. S11. A.** The mean residual fitness effect (*s* in E2 – *s* in GC) increases with background fitness (p=0.012). This implies that the fitness-mediated patterns we observe in E2 are slightly stronger in exponential phase only. **B.** Fitness effects during exponential growth correlate well with the fitness effects measured in batch culture in the E2 experiment (upper left). Correcting for measured lag effects improves the correlation (upper right), but adding saturation effects does not (lower left). Simply averaging the fitness effect measured during lag and during exponential provides the best correlation (lower right), suggesting that some of the lag correction may result from better measurement of the fitness effect during exponential phase. **C.** We detected significant differences between the fitness effects measured in E2 and the fitness effects measured during exponential phase of the growth curve experiment for one mutation in the gene MPC2 (vertical lines in the left plot show these differences for each segregant). The "lag effect" explains this deviation (center plot) but the "saturation effect" does not (right plot), suggesting that the fitness effect of this mutations involves changes in lag phase or survival during saturation phase (see "Growth curve experiment results").

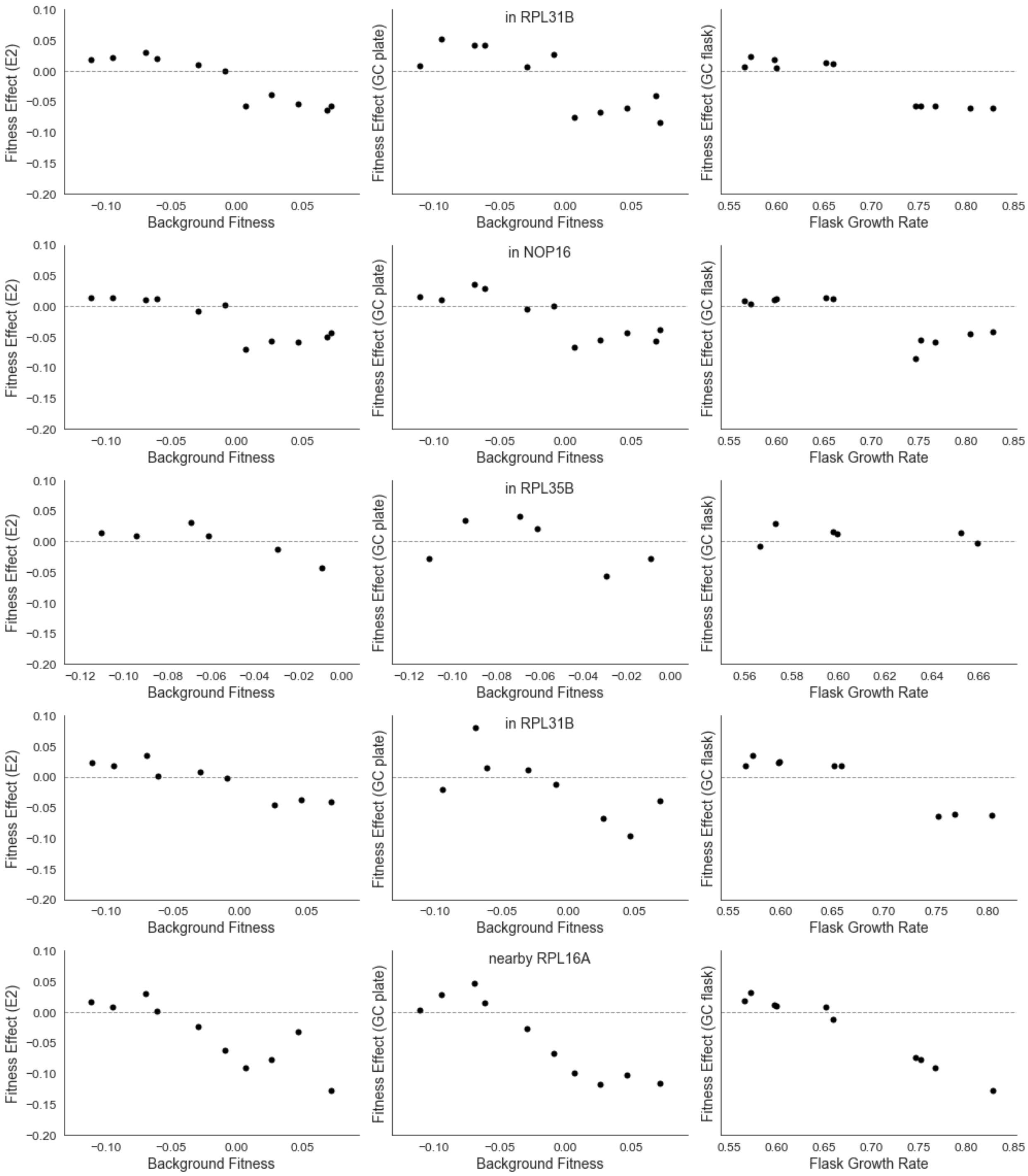

**Fig. S12.** Examples of genetic determinant plots showing that patterns from E2 (left column) are preserved when considering fitness effects during exponential growth in plates (center column) or when considering fitness effects during exponential growth in a shaken flask plotted against the exponential growth rate in a shaken flask.

### Tables

| Group | GO term | P-Value (uncorrected) | P-Value (corrected) | Associated Genes |
|---|---|---|---|---|
| qtl_chr05_360000_400000 | integral component of membrane | 0.01248853 | 0.98194444 | NCP1;KRE6;NUP188;OXA1 |
| qtl_chr05_360000_400000 | membrane | 0.01537655 | 0.98194444 | NUP188;NCP1;KRE6;OXA1;VPS36 |
| qtl_chr05_360000_400000 | ubiquitin-protein transferase activity | 0.044703 | 0.98194444 | NOT3;SLX8 |
| qtl_chr08_60000_140000 | establishment of protein-containing complex localization to telomere | 0.00234742 | 0.16934943 | SIR4;SIR3 |
| qtl_chr08_60000_140000 | negative regulation of chromatin silencing involved in replicative cell aging | 0.00234742 | 0.16934943 | SIR4;SIR3 |
| qtl_chr08_60000_140000 | telomere tethering at nuclear periphery | 0.00234742 | 0.16934943 | SIR4;SIR3 |
| qtl_chr08_60000_140000 | nuclear telomeric heterochromatin | 0.00234742 | 0.16934943 | SIR4;SIR3 |
| qtl_chr08_60000_140000 | chromatin silencing complex | 0.00234742 | 0.16934943 | SIR4;SIR3 |
| qtl_chr08_60000_140000 | nuclear chromosome, telomeric region | 0.00234742 | 0.16934943 | SIR4;SIR3 |
| qtl_chr08_60000_140000 | nucleosome binding | 0.00234742 | 0.16934943 | SIR4;SIR3 |
| qtl_chr08_60000_140000 | chromatin silencing at telomere | 0.00690812 | 0.38762203 | SIR4;SIR3 |
| qtl_chr08_60000_140000 | double-stranded DNA binding | 0.00690812 | 0.38762203 | SIR4;SIR3 |
| qtl_chr08_60000_140000 | double-strand break repair via nonhomologous end joining | 0.01355087 | 0.62210814 | SIR4;SIR3 |
| qtl_chr12_640000_720000 | lipid metabolic process | 0.01693494 | 0.95643939 | NCP1;CSR1 |
| qtl_chr14_360000_420000 | translation | 0.02526864 | 0.92963148 | RPL35B;RPS14B;RPL2A;RPL26B;RPL16A;RPL31B |
| qtl_chr14_360000_420000 | cytoplasmic translation | 0.02526864 | 0.92963148 | RPL35B;RPS14B;RPL2A;RPL26B;RPL16A;RPL31B |
| qtl_chr14_360000_420000 | ribosome | 0.02526864 | 0.92963148 | RPL35B;RPS14B;RPL2A;RPL26B;RPL16A;RPL31B |
| qtl_chr14_360000_420000 | ribonucleoprotein complex | 0.02839892 | 0.92963148 | RPL35B;RPS14B;RPL2A;NOP16;RPL26B;RPL16A;UTP22;RPL31B |
| qtl_chr14_360000_420000 | structural constituent of ribosome | 0.02526864 | 0.92963148 | RPL35B;RPS14B;RPL2A;RPL26B;RPL16A;RPL31B |
| qtl_chr14_430000_500000 | double-stranded DNA binding | 0.0435949 | 1 | DPB4;SIR4;SIR3 |
| qtl_chr15_100000_200000 | cytoplasm | 0.03954702 | 1 | GPP1;RPL2A;SFP1;PDE2;RPL16A;KAP123 |
| Top_x_effects | translation | 0.00313732 | 0.31686967 | RPL2A;RPL16A;RPL35B |
| Top_x_effects | cytoplasmic translation | 0.00313732 | 0.31686967 | RPL2A;RPL16A;RPL35B |
| Top_x_effects | cytosolic large ribosomal subunit | 0.00160425 | 0.31686967 | RPL2A;RPL16A;RPL35B |
| Top_x_effects | ribosome | 0.00313732 | 0.31686967 | RPL2A;RPL16A;RPL35B |
| Top_x_effects | structural constituent of ribosome | 0.00313732 | 0.31686967 | RPL2A;RPL16A;RPL35B |

**Table S1.** Go-term enrichments detected for genes with fitness effects influenced by multi-hit QTLs or the group of genes corresponding to the mutations with the largest background fitness coefficients in the full model. Detected enrichments only involving one gene are excluded.

| Experiment | Segregant | Replicate | Reason for Exclusion |
|---|---|---|---|
| E1 | LK1-B05 | Both | High rate of outliers |
| E1 | LK3-C01 | Both | High rate of outliers |
| E2 | LK1-A10 | Both | High Rate of Outliers |
| E2 | LK1-E07 | Both | High Rate of Outliers |
| E2 | LK1-H02 | Both | Drug resistance discrepancy in screen |
| E2 | LK1-H09 | Both | Drug resistance discrepancy in screen |
| E2 | LK2-F05 | Both | Insufficient reference/neutral barcodes |
| E2 | LK2-G09 | Both | High Rate of Outliers |
| E2 | LK3-A12 | Both | High Rate of Outliers |
| E2 | LK3-C01 | Both | High Rate of Outliers |
| E2 | LK3-B04 | Both | Drug resistance discrepancy in screen |
| E2 | LK4-B04 | Both | High Rate of Outliers |
| E2 | LK4-C07 | Both | Drug resistance discrepancy in screen |
| E2 | LK4-D12 | Both | High Rate of Outliers |
| E2 | LK6-E11 | Both | High Rate of Outliers |
| E2 | LK3-E09 | Both | Insufficient reference/neutral barcodes |
| E2 | LK2-F05 | 2 | Insufficient reference/neutral barcodes |
| E2 | LK2-F11 | 1 | Insufficient reference/neutral barcodes |
| E2 | LK2-G07 | 1 | Insufficient reference/neutral barcodes |
| E2 | LK3-A05 | 1 | Insufficient reference/neutral barcodes |
| E2 | LK3-C06 | 1 | Insufficient reference/neutral barcodes |
| E2 | LK3-C08 | 1 | Insufficient reference/neutral barcodes |
| E2 | LK3-D01 | 1 | Low sequencing coverage |
| E2 | LK3-D05 | 1 | Insufficient reference/neutral barcodes |
| E2 | LK3-G10 | 1 | Insufficient reference/neutral barcodes |
| E2 | LK3-G11 | 1 | Insufficient reference/neutral barcodes |
| E2 | LK3-H03 | 1 | Insufficient reference/neutral barcodes |
| E2 | LK4-B05 | 1 | Insufficient reference/neutral barcodes |
| E2 | LK4-E02 | 1 | Insufficient reference/neutral barcodes |
| E2 | LK4-G05 | 1 | Low sequencing coverage |
| E2 | LK5-A04 | 1 | Insufficient reference/neutral barcodes |
| E2 | LK5-E09 | 1 | Low sequencing coverage |
| E2 | LK5-F10 | 1 | Low sequencing coverage |
| E2 | LK1-D02 | 1 | Low sequencing coverage |
| E2 | LK5-H02 | 1 | Insufficient reference/neutral barcodes |
| E2 | LK5-H07 | 1 | Low sequencing coverage |
| E2 | LK6-A10 | 1 | Insufficient reference/neutral barcodes |
| E2 | LK6-F03 | 1 | Insufficient reference/neutral barcodes |
| E2 | LK1-D08 | 1 | Insufficient reference/neutral barcodes |
| E2 | LK1-D10 | 2 | Insufficient reference/neutral barcodes |
| E2 | LK1-E01 | 1 | Insufficient reference/neutral barcodes |
| E2 | LK1-G08 | 1 | Insufficient reference/neutral barcodes |
| E2 | LK1-H04 | 1 | Insufficient reference/neutral barcodes |
| E2 | LK2-A11 | 1 | Insufficient reference/neutral barcodes |
| E2 | LK2-C05 | 1 | Insufficient reference/neutral barcodes |

**Table S2.** Assays excluded from analysis.

| Segregant | Background Fitness | Excluded |
|---|---|---|
| LK1-G09 | -0.111 | No |
| LK1-B04 | -0.094 | No |
| LK3-D09 | -0.077 | Yes, was not saturated before the first transfer |
| LK5-B01 | -0.069 | No |
| LK4-B01 | -0.061 | No |
| LK2-A12 | -0.030 | No |
| LK2-D07 | -0.009 | No |
| LK6-A05 | 0.006 | No |
| LK1-G11 | 0.026 | No |
| LK1-F05 | 0.047 | No |
| LK2-B04 | 0.069 | No |
| LK1-A05 | 0.072 | No |

**Table S3.** Segregants used in the GC experiment, with information on background fitness and whether they were excluded from final analysis.

| Experiment | Hours | Hours since dilution | Cells / mL | 96-well plates sampled | µl taken per well | mL used per pellet |
|---|---|---|---|---|---|---|
| GCF | 4 | 4 | 845441.988 | NA | NA | 200 |
| GCF | 7 | 7 | 4203822.04 | NA | NA | 100 |
| GCF | 10 | 10 | 17208912.1 | NA | NA | 33.3 |
| GCF | 13 | 13 | 78753827.2 | NA | NA | 10 |
| GCF | 16 | 16 | 234874875 | NA | NA | 2 |
| GCF | 20 | 20 | 281801802 | NA | NA | 1 |
| GCF | 24 | 24 | 355835836 | NA | NA | 1 |
| GCF | 29 | 29 | 381761762 | NA | NA | 1 |
| GCF | 36 | 36 | 408798799 | NA | NA | 1 |
| GCP | 0 | 0 | 194434434 | 2 | 30 | 1 |
| GCP | 4 | 4 | 601301.992 | 8 | 80 | 60 |
| GCP | 7 | 7 | 2794569.03 | 8 | 80 | 60 |
| GCP | 10 | 10 | 10379883.3 | 8 | 80 | 30 |
| GCP | 13 | 13 | 44191526.5 | 8 | 80 | 15 |
| GCP | 16 | 16 | 110870871 | 4 | 50 | 1.5 |
| GCP | 20 | 20 | 142762763 | 4 | 50 | 1.25 |
| GCP | 24 | 24 | 173633634 | 4 | 50 | 1 |
| GCP | 29 | 29 | 179209209 | 4 | 50 | 1 |
| GCP-D1 | 31 | 7 | 2957583.03 | 8 | 80 | 60 |
| GCP-D2 | 36 | 7 | 2861229.03 | 8 | 80 | 60 |

**Table S4.** Information on timepoint sampling for the growth curve experiment. GCF is the flask experiment, GCP is the 96-well plate experiment, GCP-D1 is the timepoint 7 hours after dilution on the normal 24-hour cycle, and GCP-D2 is the timepoint 7 hours after dilution after 29 hours of growth.

| Purpose | Name | Sequence |
|---|---|---|
| First round barcode PCR | TnF1 | TCGTCGGCAGCGTCAGATGTGTATAAGAGACAGNNNNNNNNCCGTAACGTAGGTCTCTGACG |
| | TnF2 | TCGTCGGCAGCGTCAGATGTGTATAAGAGACAGNNNNNNNNTCCGTAACGTAGGTCTCTGACG |
| | TnF3 | TCGTCGGCAGCGTCAGATGTGTATAAGAGACAGNNNNNNNNTTCCGTAACGTAGGTCTCTGACG |
| | TnF4 | TCGTCGGCAGCGTCAGATGTGTATAAGAGACAGNNNNNNNNCAGCCGTAACGTAGGTCTCTGACG |
| | TnF5 | TCGTCGGCAGCGTCAGATGTGTATAAGAGACAGNNNNNNNNTGAGCCGTAACGTAGGTCTCTGACG |
| | TnF6 | TCGTCGGCAGCGTCAGATGTGTATAAGAGACAGNNNNNNNNGATAGCCGTAACGTAGGTCTCTGACG |
| | TnF7 | TCGTCGGCAGCGTCAGATGTGTATAAGAGACAGNNNNNNNNCTTACTCCGTAACGTAGGTCTCTGACG |
| | TnF8 | TCGTCGGCAGCGTCAGATGTGTATAAGAGACAGNNNNNNNNGCAGTAACCGTAACGTAGGTCTCTGACG |
| | TnF9 | TCGTCGGCAGCGTCAGATGTGTATAAGAGACAGNNNNNNNNATCTGGTTCCGTAACGTAGGTCTCTGACG |
| | TnF10 | TCGTCGGCAGCGTCAGATGTGTATAAGAGACAGNNNNNNNNAGGAACTGTCCGTAACGTAGGTCTCTGACG |
| | TnF11 | TCGTCGGCAGCGTCAGATGTGTATAAGAGACAGNNNNNNNNCAATTGGAGTCCGTAACGTAGGTCTCTGACG |
| | TnF12 | TCGTCGGCAGCGTCAGATGTGTATAAGAGACAGNNNNNNNNGGTATGTTCAACCGTAACGTAGGTCTCTGACG |
| | TnRS1 | GTCTCGTGGGCTCGGAGATGTGTATAAGAGACAGTTGCTGATAAATCTGGAGCC |
| First round barcode association PCR | Tn_R_to_P5 | TCGTCGGCAGCGTCAGATGTGTATAAGAGACAGGCGTCAGACCCCGTAGAA |
| | Tn_R_to_P7 | GTCTCGTGGGCTCGGAGATGTGTATAAGAGACAGGCGTCAGACCCCGTAGAA |
| | pTn7_edge_ to_P7 | GTCTCGTGGGCTCGGAGATGTGTATAAGAGACAGCTTCAATGGCCGCGTCGA |
| PCR for barcoding gibson assembly | Tn_BC_Amp_ Connectorator_2 | CAAACTGGACAAAAAAGATCCGTAACGTAGGTCTCTGACGCNNNNNCANNNNNCANNNNNCANNNNNCANNNNNTTCTAC GGGGTCTGACGC |
| | tTEF_to_Tn7R | TTGAACTGAACAAAATAGATCCGTGGATGGCGGCGTTAG |
| Cloning to make pUC-Hyg and pUC-yNAT | pTEF-F | AGCTTGCCTCGTCCCC |
| | tTEF-R | TGGATGGCGGCGTTAGTATCG |
| | pUC19_to_pTE F | CGGCGGGGACGAGGCAAGCTTTCGGGGAAATGTGCG |
| | pUC19_to_tTEF | GATACTAACGCCGCCATCCAACGGTTATCCACAGAATCAGG |
| Sanger sequencing to check plasmid identities | short_Tn_R | GAGGACCGAAGGAGCTAACC |

**Table S5.** Primers used in this study.

| **Antibiotic** | **Concentration for *E. coli* selection (μg/ml)** | **Concentration for *S. cerevisiae* growth / selection (μg /ml)** |
|---|---|---|
| Kanamycin | 40 | NA |
| Ampicillin | 100 | 100 (to prevent bacterial contam.) |
| clonNat | 20 | 20 |
| Hygromycin | 200 | 300 |

**Table S6.** Antibiotic concentrations used in this study

#### Data File Descriptions

**Data File S1.** Inferred fitness effects for mutations in E1 and E2 along with mutation annotation information and the results of the modeling described in "Modeling genetic determinants of fitness effects," DFE statistics from E1 and E2 along with the results of the modeling described in "Modeling genetic determinants of DFE statistics," inferred fitness effects for mutations during different stages of growth during the GC experiment, and cell density estimates and inferred growth rates for each segregant during the GC experiment.

**Data File S2.** Inferred QTLs for individual mutations in E1, for individual mutations in E2, and for DFE statistics in E2 (No QTLs were detected for DFE statistics in E1), and information on multi-hit QTL regions including whether they were observed in previous work.

**Data File S2.** Inferred fitness effects and significance testing results from our initial analysis of E1, which we used to choose mutations to include in E2.
